# Human gut microbiota interactions shape the long-term growth dynamics and evolutionary adaptations of *Clostridioides difficile*

**DOI:** 10.1101/2024.07.15.603560

**Authors:** Jordy Evan Sulaiman, Jaron Thompson, Pak Lun Kevin Cheung, Yili Qian, Jericha Mill, Isabella James, Eugenio I. Vivas, Judith Simcox, Ophelia Venturelli

## Abstract

*Clostridioides difficile* can transiently or persistently colonize the human gut, posing a risk factor for infections. This colonization is influenced by complex molecular and ecological interactions with human gut microbiota. By investigating *C. difficile* dynamics in human gut communities over hundreds of generations, we show patterns of stable coexistence, instability, or competitive exclusion. Lowering carbohydrate concentration shifted a community containing *C. difficile* and the prevalent human gut symbiont *Phocaeicola vulgatus* from competitive exclusion to coexistence, facilitated by increased cross-feeding. In this environment, *C. difficile* adapted via single-point mutations in key metabolic genes, altering its metabolic niche from proline to glucose utilization. These metabolic changes substantially impacted inter-species interactions and reduced disease severity in the mammalian gut. In sum, human gut microbiota interactions are crucial in shaping the long-term growth dynamics and evolutionary adaptations of *C. difficile*, offering key insights for developing anti-*C. difficile* strategies.

## INTRODUCTION

*Clostridioides difficile* is an opportunistic intestinal pathogen that can cause severe damage to the colon. Antibiotic treatment is highly associated with *C. difficile* infection (CDI), highlighting that microbial ecology of the human gut microbiome disrupted by antibiotics is a major determinant of *C. difficile’s* ability to colonize and induce infections^1^. *C. difficile* has been observed in up to 17% of healthy human adults ^2–6^ and up to 20-40% of hospitalized patients ^7–9^, and it can colonize individuals for 12 months or longer ^6,10,11^. Further, diarrheal events increase susceptibility to *C. difficile* and trigger long-term *C. difficile* colonization with recurrent blooms at yearlong time scales ^12^. While human gut microbiota inter-species interactions with *C. difficile* have been identified over short timescales (∼5 to 10 generations) ^13,14^, it is unclear how interactions shape the long-term growth dynamics and potential evolutionary adaptations of *C. difficile*. A deeper understanding of the ecological and molecular mechanisms shaping *C. difficile* growth over long timescales (hundreds of generations) could provide insights into mechanisms that inhibit or promote persistent colonization.

Although the impact of *C. difficile* colonization on the propensity for acquiring CDI remains unresolved ^12,15^, individuals harboring toxigenic *C. difficile* strains had a higher risk for the development of infection compared to non-colonized patients ^16,17^. The incidence of community-acquired CDI is increasing ^18^. Individuals harboring *C. difficile* could transmit it to others, serving as a reservoir for *C. difficile* ^8,19–21^. By contrast, persistent colonization of non-toxigenic strains may reduce the chance of colonization with toxigenic strains due to competition and/or stimulating a protective immune response to *C. difficile* ^22–28^. Colonization for extended periods of time can also shape *C. difficile* evolutionary adaptations ^29–31^. For example, *C. difficile* acquired a mutation in the *treR* gene that confers enhanced sensitivity to trehalose, potentially due to increased consumption of trehalose in human diets ^32^. This metabolic alteration was associated with *C. difficile*’s hypervirulence ^29^, suggesting that key evolutionary adaptations can have a major impact on pathogenic potential.

Community-level interactions have been shown to shape evolutionary adaptations ^33–35^ and lead to distinct evolutionary trajectories compared to those observed in isolation. Long-term culturing of communities constructed from the bottom up can reveal the effects of community context on the mechanisms of adaptation ^33,35–39^. This approach has been used to study model organisms ^33^ or self-assembled communities from the soil environment ^36–42^. For example, constituent community members evolved to use waste products generated by other species ^36,43,44^ and shift the community from strong competition to coexistence ^38^. Further, coevolution within a community promotes ecological diversity and stability ^39^. Increasing resource competition ^45^ or community diversity ^46^ has been shown to slow the rate of evolutionary adaptation. By contrast, positive interactions through cross-feeding promote evolutionary adaptation ^34,47,48^. Organisms that interact via metabolite exchange evolved to enhance the production of the exchanged metabolites to promote community fitness ^34^. Finally, there are many examples of species evolving new community interactions that were frequently positive^38,47,49,50^.

By building communities from the bottom up, we investigate the dynamics of *C. difficile* in various human gut communities and different environmental contexts for hundreds of generations. We demonstrate that *Eggerthella lenta* promotes stable coexistence of *C. difficile* via a growth-promoting interaction. By contrast, *P. vulgatus* or *Desulfovibrio piger* promotes instability in community dynamics across different nutrient environments. Reducing the concentration of carbohydrates in the media shifts *C. difficile* and *P. vulgatus* from competitive exclusion to coexistence by promoting metabolite exchange. In this environment, *C. difficile* adapts via point mutations in key metabolic genes. These mutations shift *C. difficile* metabolism from proline to glucose utilization, substantially alter gut microbiota inter-species interactions, and reduce disease severity in the murine gut. Further, we show that variation in nutrient landscape impacts the extent of variability of *C. difficile* growth in human gut communities. *C. difficile* abundance is lowered in an environment with a single highly accessible resource yet also displays higher variability in growth over 261 generations compared to an environment with multiple preferred carbohydrates. Overall, our study demonstrates how gut microbiota inter-species interactions influence the long-term growth dynamics and evolutionary adaptations of *C. difficile*, offering new insights for developing targeted treatments to inhibit *C. difficile*.

## RESULTS

### Human gut species differentially impact C. difficile long-term growth in communities

*C. difficile* can colonize individuals at different stages of life for variable periods of time and the factors that determine these dynamics are largely unknown. Since gut microbial ecology is a major determinant of *C. difficile* colonization, we used bottom-up microbial community experiments to investigate how interactions shape the long-term growth dynamics of *C. difficile*. To this end, we cultured *C. difficile* DSM 27147 (R20291 reference strain of the epidemic ribotype 027) with 9 diverse human gut species in a defined media (DM) that supports their growth (**Fig. 1a, S1a, Table S1-2**) ^51,52^. These human gut species are highly prevalent across individuals and span the phylogenetic diversity of the human gut microbiome ^53^. The community features *Clostridium scindens* (CS), a species previously shown to inhibit the growth of *C. difficile* in gnotobiotic mice ^54^, *Clostridium hiranonis* (CH), which can inhibit *C. difficile* via metabolic niche overlap and lead to large shifts in *C. difficile* metabolism ^55^, and *Bacteroides* species (*Bacteroides uniformis* (BU), *Bacteroides thetaoitaomicron* (BT), *P. vulgatus* (PV)), which can inhibit or promote *C. difficile* growth in different environments ^56–59^. Interactions between *D. piger* (DP), *E. lenta* (EL), *Collinsella aerofaciens* (CA), and *Prevotella copri* (PC) with *C. difficile* have also been extensively characterized on short timescales (∼5-10 generations) ^14,55^. We assembled all possible pairwise (9 total) and three-member communities (36 total) containing *C. difficile* and gut bacteria. The communities were cultured for 24 h, and an aliquot was transferred to fresh media with 40X dilution for 35 passages (∼186 generations). We performed 16S rRNA sequencing to determine the relative abundance of each species and multiplied the relative abundance by the total biomass at each time point to estimate the absolute abundance of each species ^14,60^. In addition, each species was individually cultured over time mirroring the same experimental design (**Fig. S1b**).

**Figure 1.**
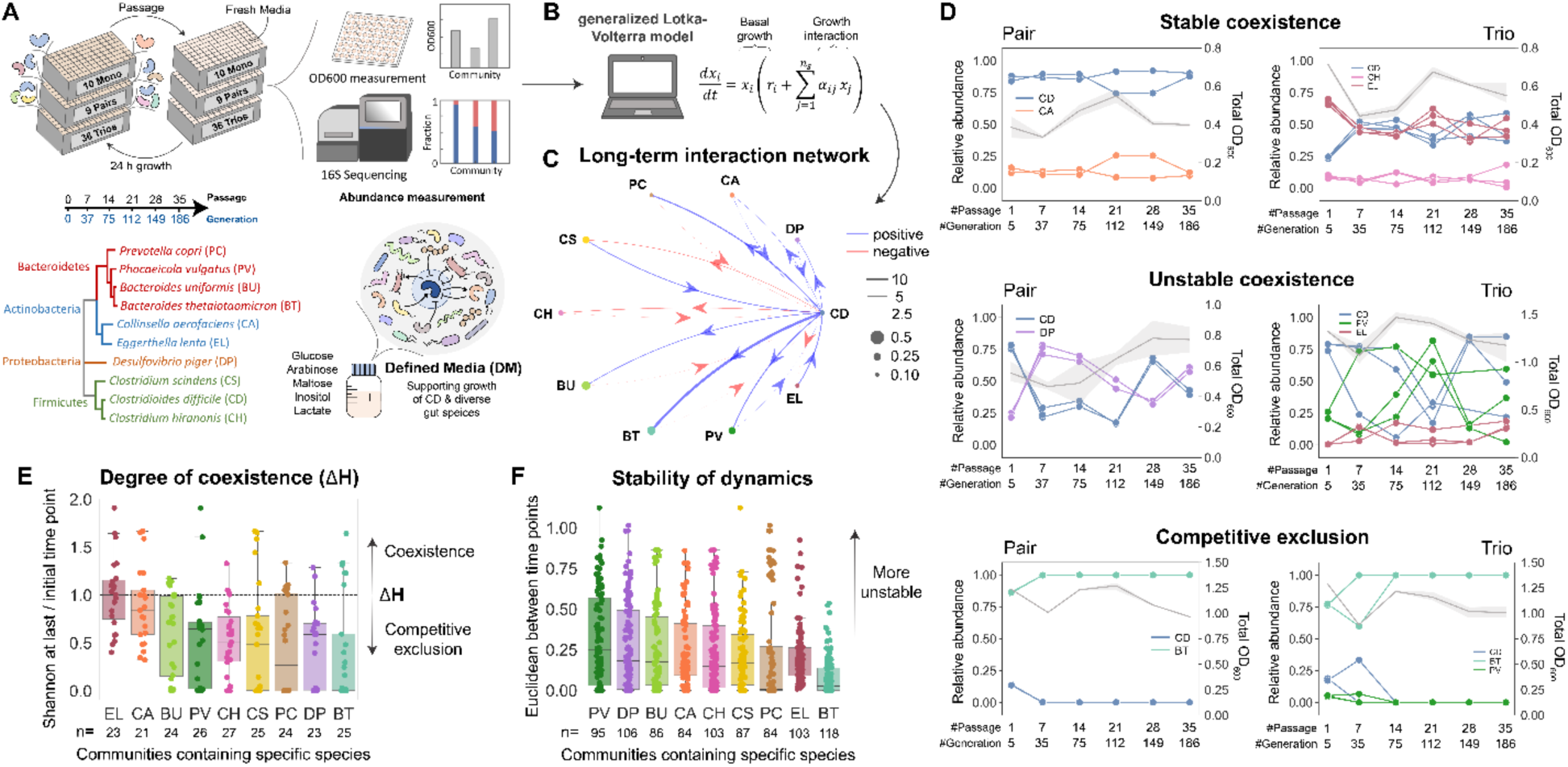
Long-term growth dynamics between *C. difficile* and human gut bacteria in pairwise and three-member communities. **a**, Schematic of the long-term growth experiment of *C. difficile* in human gut communities over 35 passages grown in the Defined Media (DM). The phylogenetic tree was generated from the 16S rRNA sequence of each species. Over time, aliquots of the cultures were subjected to OD600 measurement and multiplexed 16S rRNA sequencing to determine species abundances. **b**, Absolute abundance data from the long-term growth experiment are used to infer the parameters of a generalized Lotka–Volterra (gLV) model and elucidate the long-term interaction network of the communities (See **Methods, Table S3** DATASET001). **c**, Inferred long-term inter-species interaction network of the 9 gut species and *C. difficile* using growth data over 35 passages. Node size represents species carrying capacity and edge width represents the magnitude of the long-term inter-species interaction coefficient (aij). **d**, Representative community dynamics of *C. difficile* and gut bacteria in pairwise (left) and three-member (right) communities to show stable coexistence, unstable coexistence, and competitive exclusion. Dots connected by colored lines indicate the relative abundance of each species (left y-axis) whereas the grey lines with shaded 95% confidence interval (CI) indicate the community OD600 (right y-axis). The complete community dynamics are shown in **Fig. S1c-d**. **e**, Measurement of the degree of coexistence (⃤H) using the Shannon diversity at the last time point relative to the initial time point in all pairwise and three-member communities containing the specific species shown in the x-axis. Horizontal dashed line indicates a value of 1 (Shannon diversity at the last time point is equal to the initial time point). The complete Shannon diversity throughout the passages is available in **Fig. S4**. **f**, Box plot of Euclidean distances between time point measurements throughout 35 passages in all pairwise and three-member communities containing the specific species shown in the x-axis. For **panel e-f**, the number of data points is shown below the plots.

*C. difficile* persisted in 60% of the communities (7/9 pairwise and 20/36 three-member communities) (**Fig. S1c-d**). However, the absolute abundance of *C. difficile* decreased from the initial to the final passage in 84% of communities (**Fig. S2a**). Certain species, such as PC and BT, were enriched in communities that displayed substantial *C. difficile* inhibition. To quantify the impact of human gut species on the long-term growth of *C. difficile*, we fit a generalized Lotka–Volterra (gLV) model to the time-series data of species abundances (over 35 passages) of monoculture, pairwise, and three-member communities by simulating the passaging experimental design (**Fig. 1b**, DATASET001 in **Table S3**, **Methods**). The gLV model is a dynamic ecological model that can predict community dynamics as a function of each species’ growth and pairwise interactions with all constituent community members. This model has been used extensively to decipher inter-species interactions and predict community assembly across different environments ^14,52,60,61^. The inferred gLV inter-species interaction coefficients quantify the effect of a given species on the long-term growth of another species. The model shows good prediction performance on the measured species abundances across passages (Pearson’s R=0.89-0.91, P=3.6E-42 to 2.3E-51) (**Fig. S3a-b**).

In contrast to inferred gLV inter-species interaction networks dominated by negative interactions based on short timescales ^55^, the inferred network for long-term ecological interactions displayed a high frequency of positive interactions (**Fig. 1c**). Certain species, such as PV and DP, displayed bidirectional positive interactions with *C. difficile* over this timescale that can destabilize community dynamics based on theoretical studies ^62–64^ (**Fig. 1d, S1c-d**). By contrast, EL displayed an outgoing positive and incoming negative interaction with *C. difficile*. This topology generates a negative feedback loop that can stabilize community dynamics ^60^, providing insights into the observed stable coexistence between EL and *C. difficile* (**Fig. S1c**). Previous studies showed potential inhibitory effects of CH and CS on *C. difficile* ^14,55,65–69^, consistent with the inhibition of *C. difficile* by CH inferred by the model. However, CH and CS were outcompeted by *C. difficile* in pairwise co-culture after 7 passages and were frequently excluded from the 3-member communities (**Fig. S1c-d, S2b**). Therefore, interventions that enhance the abundance of CH and CS in human gut communities may be critical to achieving robust inhibition of *C. difficile* over long timescales.

To evaluate the degree of coexistence in the community, we calculated the change in Shannon diversity over time (⃤H) (**Fig. 1e, S4**). Communities containing EL displayed the highest ⃤H, whereas communities containing BT displayed the lowest ⃤H. In addition, we quantified the extent of variability in community dynamics using the Euclidean distance of species’ relative abundances between each pair of time points. Large values of Euclidean distance indicate community instability, whereas small values indicate temporal stability. The presence of PV and DP yielded larger Euclidean distances than other species (**Fig. 1f**), consistent with their bidirectional positive interaction network topology.

We investigated community dynamics of a subset of communities that displayed coexistence with *C. difficile* over longer timescales and across different environmental conditions. Specifically, we cultured pairwise communities containing CA, EL, BU, DP, and PV with two distinct *C. difficile* strains in three different media conditions for 341 generations (**Fig. S5a-d**). Since *C. difficile* has large genetic variability across strains ^70–73^, we characterized a toxigenic DSM 27147 strain and a non-toxigenic MS001 strain with a larger genome and distinct metabolic capabilities ^55^. In addition, we analyzed three nutrient environments: DM, DM with a 4-fold lower carbohydrate concentration, and DM with a 2-fold higher amino acid concentration. Decreasing the concentration of carbohydrates could influence *C. difficile* growth by altering the strength of competition and/or media acidification by certain gut species. Increasing amino acid availability could enhance the growth of *C. difficile* due to Stickland metabolism ^74^. Overall, *C. difficile* DSM and MS001 displayed similar long-term dynamics across communities and nutrient environments except in co-culture with PV in DM and DM with increased amino acids, and with DP in DM with limited carbohydrates (**Fig. S5e-g**). The dynamics of CD-PV and CD-DP exhibited larger instability (**Fig. S5h-j**) and displayed larger variation across the distinct nutrient environments than other communities (**Fig. S5e-g**).

The long-term growth dynamics of *C. difficile* with PV or DP displayed qualitative differences across different experiments when cultured via serial dilutions in the same media (DM) (**Fig. S1c, S5e**). For instance, while species relative abundance alternated between high and low values across 35 passages in the CD-PV community in one experiment (**Fig. S1c**), *C. difficile* was excluded from the community between the 35^th^ to 42^nd^ passage in a different experiment (**Fig. S5e**). This suggests that the observed instability in community dynamics yielded an elevated risk of extinction, consistent with previous theoretical studies ^45^. This variability in growth dynamics was also observed across biological replicates containing CD and PV (e.g. in CD-PV-CS), where one replicate displayed unstable coexistence and another displayed competitive exclusion.

In sum, certain species such as EL promote stable coexistence of *C. difficile* in communities across long timescales. By contrast, PV or DP promotes instability in community dynamics across different environmental conditions. These dynamics are overall consistent with their inferred interaction network topologies.

### Resource-limited environment promotes metabolite cross-feeding and coexistence between C. difficile and P. vulgatus

Previous studies have shown that 30% of bacteria in the human colon are *Bacteroides* species, with PV, BT, BU*, B. distasonis, B. fragilis*, and *B. ovatus* displaying the highest prevalence across individuals ^75–78^. Studying interactions between *C. difficile* and these highly abundant and prevalent species could provide insights into the factors shaping *C. difficile* colonization. CD-PV co-culture displays a unique feature where modifying the nutrient environment could alter community dynamics from competitive exclusion to coexistence over this timeframe (**Fig. S5e-f**). In the presence of high carbohydrate concentrations, *C. difficile* was excluded between ∼186-224 generations, whereas in media with reduced carbohydrate concentration, *C. difficile* coexisted with PV over 341 generations.

To uncover the metabolic activities driving coexistence versus competitive exclusion in these conditions, we performed exo-metabolomic profiling on PV and *C. difficile* in both media (**Fig. 2a-b**). In the limited carbohydrate media, our results were consistent with the cross-feeding of multiple metabolites from PV to *C. difficile* (**Fig. 2a, S6**). In monoculture, *C. difficile* consumed many amino acids in the media due to its ability to perform Stickland metabolism ^65,74,79,80^. Notably, PV released 84% of the metabolites that *C. difficile* utilized, including amino acids for Stickland metabolism. These metabolites displayed a larger decrease in abundance in the CD-PV co-culture than in the PV monoculture, suggesting that they are being consumed by *C. difficile*. The release of amino acids and cross-feeding from *Bacteroides* to *Clostridium* species has been previously reported ^81,82^. By contrast, the predicted cross-feeding is sparse in the high carbohydrate media where 12% of the metabolites that *C. difficile* utilized were released by PV (**Fig. 2b, S7**). This is consistent with a previous study where high concentrations of acetate suppressed the release of many metabolites from *Bacteroides* including amino acids ^83^. Further, 24% of the metabolites that *C. difficile* consumed are also consumed by PV in monoculture, implying a higher degree of resource competition in this environment. In this media, both *C. difficile* and PV utilized glucose and inositol. Of the metabolites that *C. difficile* consumed in the high carbohydrate media, 45% were also consumed in the limited carbohydrate media.

**Figure 2.**
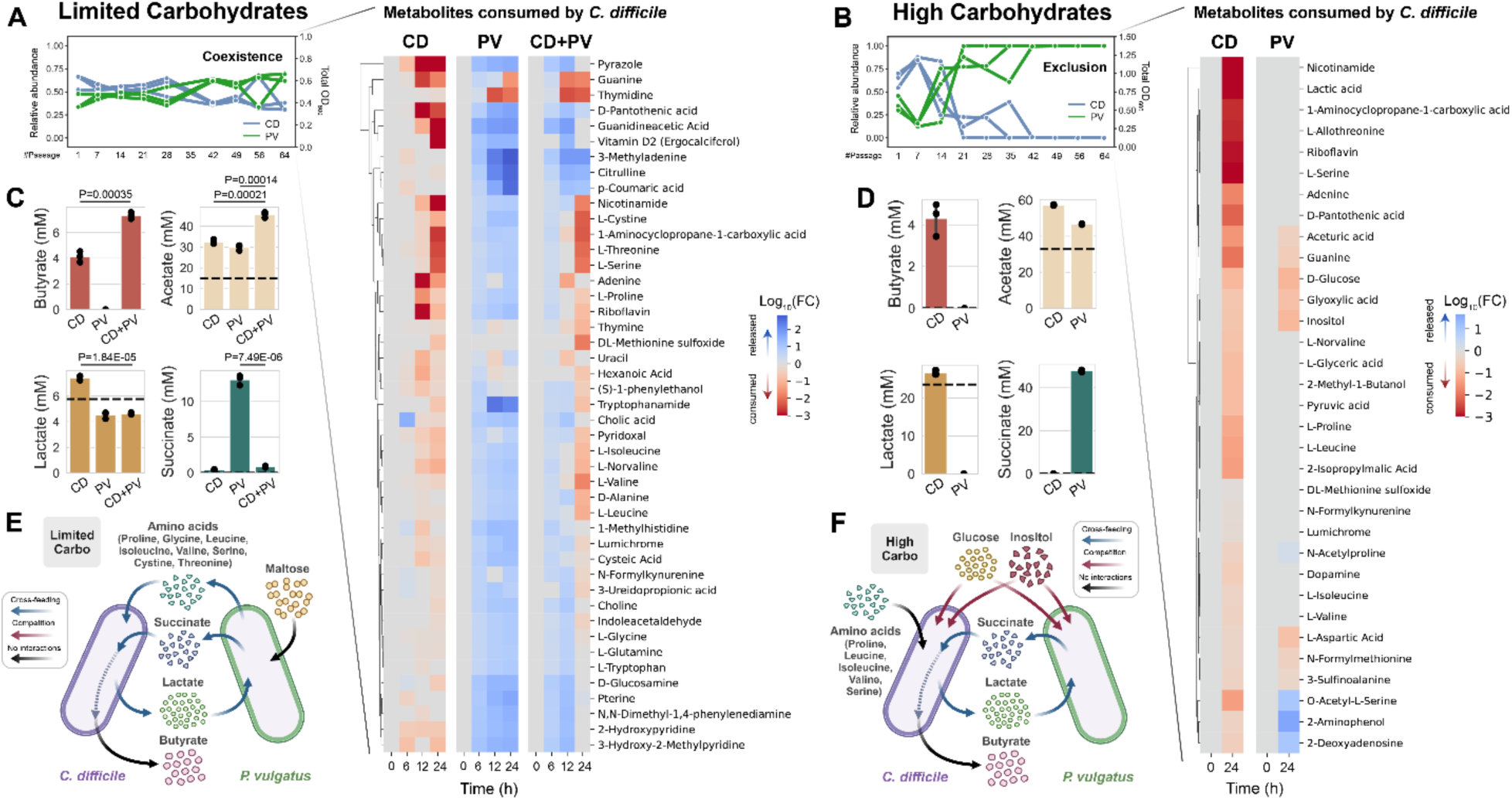
Exo-metabolomic profiling of *C. difficile* and PV in two media conditions. **a-b**, Heatmap of the fold change of metabolites consumed by *C. difficile* in DM with reduced carbohydrate concentration (limited carbohydrate media) (**a**) or DM (high carbohydrate media) (**b**) compared to the blank media (t = 0 h). The complete metabolomics profile is shown in **Fig. S6** for DM with limited carbohydrates and **Fig. S7** for DM. The figures of the time-course abundance measurement were taken from **Fig. S5e** and **f**. **c-d**, Quantification of organic acids in DM with reduced carbohydrate concentration (**c**) and DM (**d**) (mean ± s.d., n=3). Horizontal dashed lines indicate the concentration detected in the blank media. The *p*-values from unpaired *t*-test (two-sided) are shown. **e-f**, Schematic of major metabolic activities in CD-PV co-culture in DM with reduced carbohydrate concentration (**e**) and DM (**f**). Parts of the figure are generated using Biorender.

By performing exo-metabolomics on the other 8 human gut species in monocultures, our results suggested that potential cross-feeding is unique to the PV-CD community in the limited carbohydrate media. Specifically, only ∼3-15% of the metabolites released by other gut species were utilized by *C. difficile* (**Fig. S7a-c, f**). Of the 9 gut bacteria, CS and CH have the highest metabolite utilization overlap with *C. difficile* and could compete with *C. difficile* over Stickland amino acids ^65,84^. This potential resource competition is consistent with their extinction in co-culture with *C. difficile* (**Fig. S1c**). CS and CH also have lower growth rates than *C. difficile* (**Fig. S8a-c**). In addition to cross-feeding, the coexistence of species can be mediated via the utilization of unique resources ^85^. Of all metabolites consumed by *C. difficile* in the high carbohydrate media, 21% are unique (**Fig. S7f**). This implies that orthogonal niches may enable the coexistence of *C. difficile* with certain gut species in this media, such as CA, BU, and EL (only 9%, 15%, and 21% metabolite utilization overlap respectively) (**Fig. 1e, S1c-d**). The number of metabolites consumed by *C. difficile* was the second largest after CS (**Fig. S7d**), suggesting that *C. difficile* has a flexible metabolic niche.

Fermentation end products play key roles in inter-species interactions in the human gut microbiome ^86^. Therefore, we quantified the concentration of butyrate, lactate, acetate, and succinate in the supernatants of *C. difficile* and PV after 24 h of growth (**Fig. 2c-d**). While acetate was produced by *C. difficile* and PV, PV produced succinate, and *C. difficile* produced butyrate in both media conditions. In co-culture, succinate produced by PV was substantially reduced, and butyrate produced by *C. difficile* was higher than in monoculture (**Fig. 2c**). This implies that *C. difficile* used succinate released by PV to produce butyrate ^74^. Cross-feeding of succinate from *Bacteroides* to *C. difficile* has also been observed in mice, suggesting that this metabolic exchange is relevant for the mammalian gut ^56^. In addition, *C. difficile* produced lactate that was consumed by PV in monoculture and co-culture (**Fig. 2c-d**). In sum, these data suggest that PV and *C. difficile* can interact via metabolite exchange of multiple metabolites.

Overall, an environment with limited carbohydrates promotes metabolite cross-feeding and coexistence between *C. difficile* and PV (**Fig. 2e**). In the presence of high carbohydrate concentrations, resource competition may dominate over cross-feeding, thus promoting the exclusion of *C. difficile* in co-culture with PV after ∼35-42 passages (**Fig. 2f**). In addition, *C. difficile* uniquely consumes a subset of metabolites in this media, which could contribute to the observed coexistence with other human gut species such as CA and EL.

### Evolutionary adaptations of C. difficile in co-culture with PV leads to altered metabolic activities

Prolonged coexistence between *C. difficile* and PV could lead to evolutionary adaptations in *C. difficile*. In the limited carbohydrate media, *C. difficile* abundance decreased over time (**Fig. S5f, 2a**). To determine if the change in abundance stemmed from an evolutionary adaptation, we isolated *C. difficile* from the final passage of the CD-PV community. The isolated *C. difficile* strain (evolved strain) displayed a lower relative abundance than the ancestral strain in co-culture with PV in the media used to isolate the strain (**Fig. S9a**). Notably, when co-cultured with PV over 341 generations in the limited carbohydrate media, the evolved *C. difficile* strain displayed a converging trend towards approximately equal species proportion, whereas the abundance of the ancestral strain was diverging away from equal proportions (**Fig. S9b-c**). This implies that the evolved *C. difficile* strain could coexist better with PV than the ancestral strain over this timescale.

Whole-genome sequencing (WGS) revealed that the isolated *C. difficile* strain harbored two single-point non-synonymous mutations (**Fig. 3a, Table S4**). The mutations are located in *gene 206* (G533W) expressing the phosphotransferase system (PTS) sugar transporter subunit IIA, and in *prdR* (A341V), a central metabolism regulator that controls preferential utilization of proline and glycine to produce energy via the Stickland reactions. In the presence of proline, PrdR activates transcription of the proline reductase-encoding genes and negatively regulates the glycine reductase-encoding genes ^79^. Sanger sequencing of the *prdR* gene confirmed the presence of A341V mutation in *C. difficile* isolated from two other biological replicates of the CD-PV pair. This mutation was not present in other pairwise communities (CD-BU, CD-CA, CD-DP, CD-EL) or *C. difficile* that was passaged alone in the same media condition, suggesting that the mutation uniquely emerged in co-culture with PV. By subjecting the whole CD-PV population to WGS, we also detected the mutation in *gene 206* (G533W) in *C. difficile* across all three biological replicates of the CD-PV pairwise community. In addition, we identified several mutations arising in PV’s genome in the CD-PV pairwise community that did not reach fixation by the 64^th^ passage (∼18 to 46% of the population) (**Table S5**). Notably, some of the genes with identified non-synonymous mutations were frequently mutated in *Bacteroides* in healthy humans ^87^.

**Figure 3.**
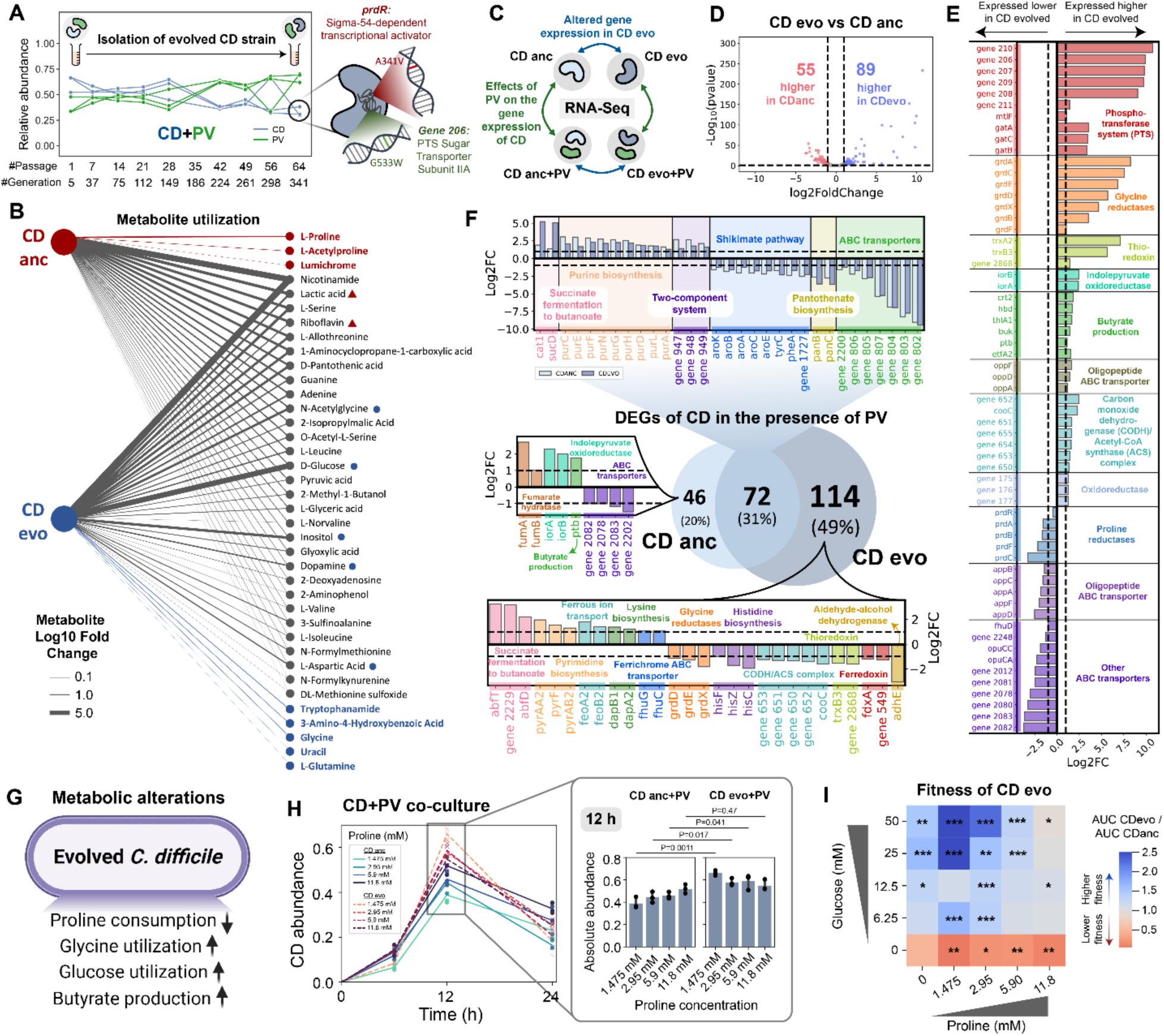
*C. difficile* undergoes evolutionary adaptations that alter its metabolism after prolonged co-culture with *P. vulgatus*. **a**, *C. difficile* was isolated from CD-PV grown in limited carbohydrate media after 341 generations. Figure of the community dynamics was taken from **Fig. S5f**. **b**, Bipartite network of metabolite utilization between the ancestral and evolved *C. difficile* strains in DM. Metabolites shown have significantly lower concentrations than the blank media (two-sided t-test with unequal variance). Metabolites bolded with red (blue) are uniquely utilized by the ancestral strain (evolved strain). Metabolites marked with red triangle (blue circle) asterisks have >10-fold higher utilization in the ancestral strain (evolved strain). **c**, Schematic of the genome-wide transcriptional profiling experiment. **d**, Volcano plot of log-transformed transcriptional fold changes of the evolved *C. difficile* strain compared to the ancestral strain. Vertical dashed lines indicate 2-fold change and horizontal dashed line indicates the statistical significance threshold as calculated by DESeq2’s Wald test with Benjamini-Hochberg multiple testing correction (BH-adjusted p=0.05). **e**, Log-transformed fold changes of selected genes that were expressed higher and lower in the evolved *C. difficile* strain compared to the ancestral strain. Vertical dashed lines indicate a 2-fold change. **f**, Comparison of DEGs in the ancestral and evolved *C. difficile* strains in the presence of PV. **g**, Schematic of metabolic alterations in the evolved *C. difficile* strain. **h**, *C. difficile* abundance when grown with PV under different proline concentrations. Solid (dashed) lines indicate ancestral (evolved) *C. difficile* strain when co-cultured with PV. Bar plots show *C. difficile* abundance at 12 h (mean ± s.d., n=3). *p*-values from unpaired *t*-test (two-sided) between evolved and ancestral strains are shown. **i**, Heatmap of fitness of the evolved *C. difficile* strain compared to the ancestral strain in response to varying glucose and proline concentrations. Fitness comparison is quantified by the Area Under the Curve (AUC) ratio based on monoculture growth (**Fig. S14**). Asterisks indicate *p*-value from unpaired *t*-test between evolved and ancestral strains. *** indicates p<0.001, ** indicates p<0.01, * indicates p<0.05, ns indicates not significant. Exact p-values are shown in **Fig. S14**. Parts of the figure are generated using Biorender.

Since these mutations could impact metabolic activities, we characterized the difference in exo-metabolomic profile of the evolved and ancestral *C. difficile* strains in the high carbohydrate media (DM) (**Fig. 3b, S10**) and limited carbohydrate media (**Fig. S6**). While the ancestral strain consumed proline consistent with previous knowledge of its metabolism ^55,79^, proline in DM was not significantly reduced by the evolved strain (**Fig. 3b**). By contrast, the consumption of glycine, aspartic acid, and N-acetylglycine (a derivative of glycine) was enhanced by the evolved strain. Similarly, in the limited carbohydrate media, the evolved *C. difficile* strain displayed a reduced rate of proline utilization compared to the ancestral strain both in monoculture and co-culture with PV (**Fig. S6d**). By contrast, the consumption rate of many other amino acids including glycine, leucine, isoleucine, valine, and serine was higher in the evolved *C. difficile* strain both in monoculture and co-culture with PV. In addition to amino acids, the evolved strain consumed more glucose than the ancestral strain in both media conditions.

To provide insights into how the mutations affect the gene expression of *C. difficile,* we performed genome-wide transcriptional profiling on the ancestral and evolved *C. difficile* strains in the absence and presence of PV (**Fig. 3c, S11a-b**). Since one of the mutations plays a role in carbohydrate utilization, we cultured cells in DM where *C. difficile* consumed glucose and inositol in addition to amino acids (**Fig. 2f**). Of 3,508 total genes, 89 and 55 displayed significantly higher and lower expression respectively in the evolved *C. difficile* than the ancestral strain in monoculture (**Fig. 3d-e, Table S6**). PTS-related genes, including the mutated gene 206, were expressed substantially higher in the evolved strain. This suggests that the evolved strain adapted to transport and phosphorylate carbohydrates more effectively than the ancestral strain, contributing to the observed higher utilization of carbohydrates such as glucose (**Fig. 3b**). Although the change in *prdR* expression was moderate (1.4-fold up-regulated), there was a massive shift in the expression of the *prd* and *grd* operon. Specifically, the glycine reductase-encoding genes were highly up-regulated (2.1-305-fold), whereas the proline reductase-encoding genes were down-regulated (3.0-9.7-fold) in the evolved strain compared to the ancestral (**Fig. 3e**). This is consistent with the alterations in proline and glycine utilization in the evolved strain as observed through exo-metabolomic profiling (**Fig. 3b, S6d**).

Other metabolic genes such as the carbon monoxide dehydrogenase/acetyl-CoA synthase complex that is responsible for the carbonyl branch of the Wood–Ljungdahl Pathway (WLP), converting CO2 to acetyl-CoA, were expressed higher in the evolved strain compared to the ancestral (**Fig. 3e**). In the evolved strain, genes for butyrate production including *thlA1, hbd, crt2* (converting acetyl-CoA to butyryl-CoA) and *ptb* and *buk* (converting butyryl-CoA to butyrate), were also expressed higher. The evolved strain displayed 2.7-fold higher butyrate production than the ancestral strain (**Fig. S12a**). This higher level of butyrate produced by the evolved *C. difficile* strain (11.6 mM) is comparable to a major butyrate-producing bacteria *Coprococcus comes* cultured in a similar media ^51,88^ and could potentially influence disease severity *in vivo* ^89^. Previous studies have shown that certain *Bacteroides* species displayed nutrient-specific growth sensitivity towards butyrate ^90^. Of the characterized *Bacteroides* species, only PV displayed significant growth reduction under intermediate butyrate concentrations. The butyrate concentration produced by the evolved strain was in the lower-inhibitory regime of the butyrate dose-response curve for PV (**Fig. S12b**).

In co-culture with PV, both the ancestral and evolved *C. difficile* strains up-regulated genes for succinate fermentation to butanoate, with the evolved strain exhibiting higher expression of these genes (**Fig. 3f**, **S11c-f, Table S7-9**). Since the growth media does not contain succinate, this implies cross-feeding from PV, consistent with the organic acids measurements data (**Fig. 2c-d**). The gene expression profile of PV also displayed substantial differences in co-culture with the ancestral and evolved *C. difficile* strain. Notably, PV exhibited higher expression of many genes involved in amino acid biosynthesis in the presence of the evolved *C. difficile* strain compared to the ancestral strain (**Fig. S11g**). This indicates that PV’s amino acid biosynthesis is either induced by the evolved *C. difficile* strain or inhibited by the ancestral strain, which might contribute to the observed differences in the long-term growth dynamics between the ancestral and evolved *C. difficile* strain with PV (**Fig. S9b**). Since the evolved *C. difficile* strain has a higher consumption of many amino acids (**Fig. S6d**), this implies an enhanced strength of amino acid cross-feeding with PV, providing insights into a mechanism that may enhance stable coexistence with PV (**Fig. S9b**).

Overall, evolutionary adaptations that altered *C. difficile* metabolism (**Fig. 3g**) and increased its ability to coexist with PV (**Fig. S9b**) arose after prolonged coexistence with PV.

### Growth of evolved C. difficile strain is less limited by proline and enhanced in the presence of high glucose concentrations

A key unresolved question is how this metabolic adaptation impacts the fitness of *C. difficile* across different environments. To evaluate changes in fitness, we characterized the growth of the evolved and ancestral *C. difficile* strains in the presence of different amino acid concentrations. The evolved *C. difficile* strain displayed higher growth than the ancestral strain in the presence of reduced amino acid concentrations (20%) at earlier time points, whereas the growth responses of the two *C. difficile* strains were similar in other conditions (**Fig. S9d**). The EC50 of the evolved *C. difficile* strain for proline (concentration of proline that yields 50% of the maximum growth) was substantially lower than the ancestral strain (**Fig. S13a, e**). This indicates that the evolved strain can compete more efficiently for low proline concentrations than the ancestral strain. Proline is the growth-limiting resource for the ancestral *C. difficile* strain in our media, which has been observed in many other *C. difficile* clinical isolates with diverse genomes ^55^. However, proline is not the limiting substrate for the evolved strain as evidenced by our metabolomics data (**Fig. 3b**). Consistent with this result, the abundance of the evolved strain did not vary with proline in co-culture with PV, indicating that the growth of the evolved *C. difficile* strain was in the saturated regime of proline concentrations (**Fig. 3h, S13g**). By contrast, the abundance of the ancestral strain increased with proline concentration (i.e. linear regime of proline dose-response). The abundance of the evolved *C. difficile* strain was higher than the ancestral strain in co-culture with PV at 12 h under low initial proline concentration and then decreased to the final time point (**Fig. 3h**). The evolved and ancestral strains displayed similar growth as a function of glycine in monoculture (**Fig. S13b, d**) and in co-culture with PV (**Fig. S13h**).

The glucose EC50 of the evolved *C. difficile* strain was higher than the ancestral strain (**Fig. S13f**). In addition, the evolved strain displayed higher biomass than the ancestral strain under high glucose concentrations (**Fig. S13c, f**), but similar size and morphology at the single-cell level (**Fig. S9e**). This implies that the evolved strain more efficiently utilizes high glucose concentrations. In the low-concentration regime, the evolved *C. difficile* strain displayed a trade-off between increased sensitivity towards proline and reduced utilization of glucose. In co-culture with PV, the evolved *C. difficile* strain displayed lower abundance than the ancestral strain at low glucose concentrations but displayed similar growth at high glucose concentrations (**Fig. S13i**). This is consistent with its reduced growth compared to the ancestral strain in the presence of PV in the limited carbohydrate media (**Fig. S9a**).

In sum, the shift in metabolic activities in the evolved *C. difficile* strain (**Fig. 3g**) impacts its fitness across different combinations of proline and glucose. The fitness of the evolved strain was enhanced in the presence of high glucose and low proline concentrations, whereas the ancestral strain displayed higher fitness in the absence of glucose and the presence of proline (**Fig. 3i, S14**).

### Evolved C. difficile strain displayed altered inter-species interactions with human gut bacteria

To investigate whether the observed metabolic shifts in the evolved *C. difficile* strain impact human gut microbiota inter-species interactions, we constructed 96 combinations of 2-8 member communities containing the evolved or the ancestral *C. difficile.* The other species span the phylogenetic diversity of the human gut microbiome, are highly prevalent across the human population (CS, CH, DP, BT, PV, BU, and CA), and have been extensively characterized in different media (**Fig. 4a**) ^14,55,60^. We fit the gLV model to the time-series data of species absolute abundances (0, 12, and 24 h) (**Table S3** DATASET002, **Fig. S15a-c**). To evaluate model prediction performance on held-out data, we performed 10-fold cross-validation where the model was trained on a fraction of the data and then used to evaluate prediction performance on the held-out community measurements (**Fig. S15d,** see **Methods**). Using a 10-fold cross-validation, the model prediction exhibited good agreement with the measured species abundance in all communities containing the ancestral and the evolved *C. difficile* strain (Pearson’s R=0.95-0.99, P<10E-05). However, certain species such as CH, displayed a low to moderate prediction performance ^55^. This might be due to insufficient variation of the particular species abundance across communities or limited flexibility of the gLV model to capture complex interaction modalities ^91^.

**Figure 4.**
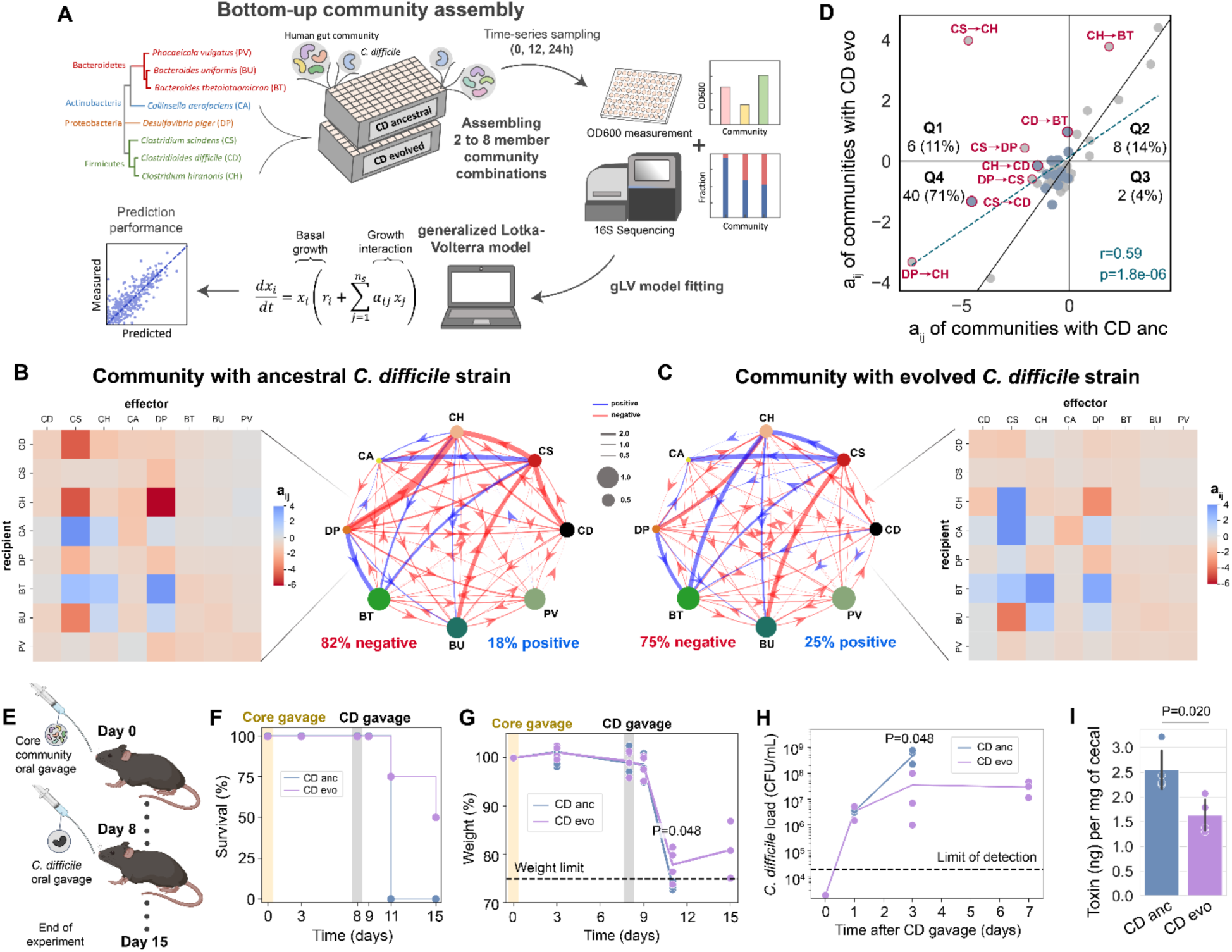
Evolved *C. difficile* strain displayed alteration in inter-species interactions with human gut bacteria and reduced disease severity in the mammalian gut. **a**, Schematic of workflow to decipher interactions between ancestral and evolved *C. difficile* strain and human gut bacteria (See **Methods, Table S3** DATASET002). The gLV model was fit to species absolute abundance and the inferred gLV parameters revealed inter-species interactions. **b-c**, Inferred inter-species interaction networks between the 7 gut species and the ancestral (**b**) and evolved (**c**) *C. difficile* strain. Node size represents species carrying capacity and edge width represents the magnitude of the inter-species interaction coefficient (aij). The heatmaps show the aij among the 8 species in the community. **d**, Scatter plot of the aij between communities containing ancestral versus evolved *C. difficile* strain. Grey data points are aij between two gut species, whereas blue data points are aij between *C. difficile* and a gut species. Blue dashed line indicates linear regression between the aij values of the two communities. Two-sided Pearson’s correlation coefficient (*r*) and *p*-values are shown. **e,** Schematic of the mice experiment. Mice were gavaged with a 7-member bacterial community for 8 days prior to challenge with the ancestral or evolved *C. difficile* strain. **f**, Percent survival of mice gavaged with ancestral and evolved *C. difficile* strain. **g**, Percent of initial weight of mice gavaged with ancestral and evolved *C. difficile* strain. Data points indicate individual mice, and the line indicates the average of all mice in the group. Horizontal dashed line indicates the weight limit of 75%. Mice with weights that dropped below the limit were sacrificed. **h**, *C. difficile* load in the fecal (survived mice) and cecal (dead mice) content as determined by CFU counting on *C. difficile* selective plates. Horizontal dashed line indicates limit of detection. For panel **g-h**, significant *p*-values from unpaired *t*-test (two-sided) between mice gavaged with ancestral vs. evolved *C. difficile* strain are shown. **i**, Toxin concentration per mg of cecal content. Data were shown as mean ± s.d. (n=4). *p*-value from unpaired *t*-test (two-sided) is shown. Parts of the figure are generated using Biorender.

Based on the inferred gLV parameters, both pairwise interaction networks displayed a high frequency of negative interactions, consistent with previous findings ^14,55^ (**Fig. 4b-c**). The frequency of positive interactions in the community was higher in the presence of the evolved (25%) versus ancestral (18%) *C. difficile* strains. In addition, 15% of inter-species interactions displayed inconsistent signs in the evolved versus ancestral inter-species interaction network (**Fig. 4d**). This implies that the two single-point mutations in the evolved strain are critical determinants of gut microbiota inter-species interactions. While CS and CH strongly inhibited the ancestral *C. difficile’*s growth, this inhibition is substantially suppressed for the evolved *C. difficile* strain. Since the evolved *C. difficile* strain uses more glucose and less proline (**Fig. 3b**), this could relieve the competition over proline with CS and CH. Beyond pairwise interactions between *C. difficile* and gut species, the evolutionary adaptations of *C. difficile* also indirectly impacted pairwise interactions between other constituent community members, mainly CS and CH with other gut species (**Fig. 4d**). Overall, *C. difficile*’s evolutionary adaptation yielded substantial direct and indirect alterations of human gut microbiota inter-species interactions. These changes could emerge due to the shifts in the metabolic niche of the evolved *C. difficile* strain (**Fig. 3, S6, S10**).

### Evolved C. difficile strain reduced disease severity in the mammalian gut

Amino acids such as proline have been shown to regulate *C. difficile* toxin production *in vitro* ^79,92–94^ and influence colonization in the mammalian gut ^80,95^. To determine if the *C. difficile* evolutionary adaptations alter toxin production, we characterized toxin expression using the Enzyme-Linked Immunosorbent Assay (ELISA) after 24 h of growth. The evolved *C. difficile* strain displayed substantially lower toxin concentration and yield (toxin concentration divided by the OD600 of *C. difficile*) compared to the ancestral strain across a wide range of amino acid concentrations (**Fig. S16**).

To examine whether the metabolic adaptations in the evolved *C. difficile* strain impact disease severity, we orally gavaged germ-free mice with the 7-member synthetic human gut community (CS, CH, DP, BT, PV, BU, and CA) for 8 days to allow time for the establishment of a stable human gut community and immune system training ^96^ (**Fig. 4e**). After 8 days, a group of mice was orally gavaged with the ancestral *C. difficile* strain, and another group was gavaged with the evolved *C. difficile* strain. Three days after *C. difficile* inoculation, all the mice gavaged with the ancestral *C. difficile* strain died (**Fig. 4f-g**). By contrast, only 50% of the mice harboring the evolved *C. difficile* strain died after 3 and 7 days of *C. difficile* colonization. The relative reduction in weight for mice harboring the evolved *C. difficile* strain was significantly lower than the mice harboring the ancestral *C. difficile* strain, although both groups displayed a decreasing trend in weight over the first few days after *C. difficile* challenge. After 3 days of *C. difficile* challenge, the mean fraction of the ancestral and evolved *C. difficile* strain in the community was 10.6% and 3.3% respectively (**Fig. S17**). Mice gavaged with the evolved *C. difficile* strain displayed significantly lower *C. difficile* abundance and toxin concentration compared to the mice gavaged with the ancestral *C. difficile* strain (**Fig. 4h-i**). In sum, the evolved *C. difficile* strain displayed reduced colonization ability and reduced disease severity in the murine gut compared to the ancestral *C. difficile* strain.

### Nutrient environments impact variability in long-term community assembly

Community dynamics have different degrees of stability over the passages (**Fig. 1f, S5h-j**), and higher instability leads to larger variability. Characterizing the growth of *C. difficile* in communities over long timescales across a larger number of replicates could provide insights into how composition diffuses over periodic regrowth cycles due to variability in growth dynamics ^41^. Understanding the factors that modulate variability in community assembly is important as it could influence the ability of *C. difficile* to persist in the community over time. To evaluate the degree of variability in *C. difficile* growth within a community after long-term passaging, *C. difficile* was introduced into an 8-member community with 96 biological replicates (**Fig. S18a**). *C. difficile* persisted in the community with a nearly constant mean of absolute abundance, but the variability in abundance across replicates substantially increased from the first to the 14^th^ passage and then increased again from the 42^nd^ to the 56^th^ passage (**Fig. S18c-e**).

The impact of environments with multiple *C. difficile*-preferred carbohydrates versus glucose as a single highly accessible resource on gut microbiota inter-species interactions with *C. difficile* has been studied ^55^, but their role in shaping the variability of *C. difficile* growth is not known. We cultured *C. difficile* with human gut communities in the presence of glucose as a single highly accessible resource or multiple *C. difficile*-preferred carbohydrates mirroring post-antibiotic environments where numerous resources could be exploited by *C. difficile* (sorbitol ^97,98^, mannitol ^97,98^, trehalose ^29,32^, and succinate ^56,97^) (**Fig. 5a**). In monoculture, all gut species consumed glucose, but sorbitol was consumed by CS and CH, and mannitol was consumed by CH (**Fig. S19a**).

**Figure 5.**
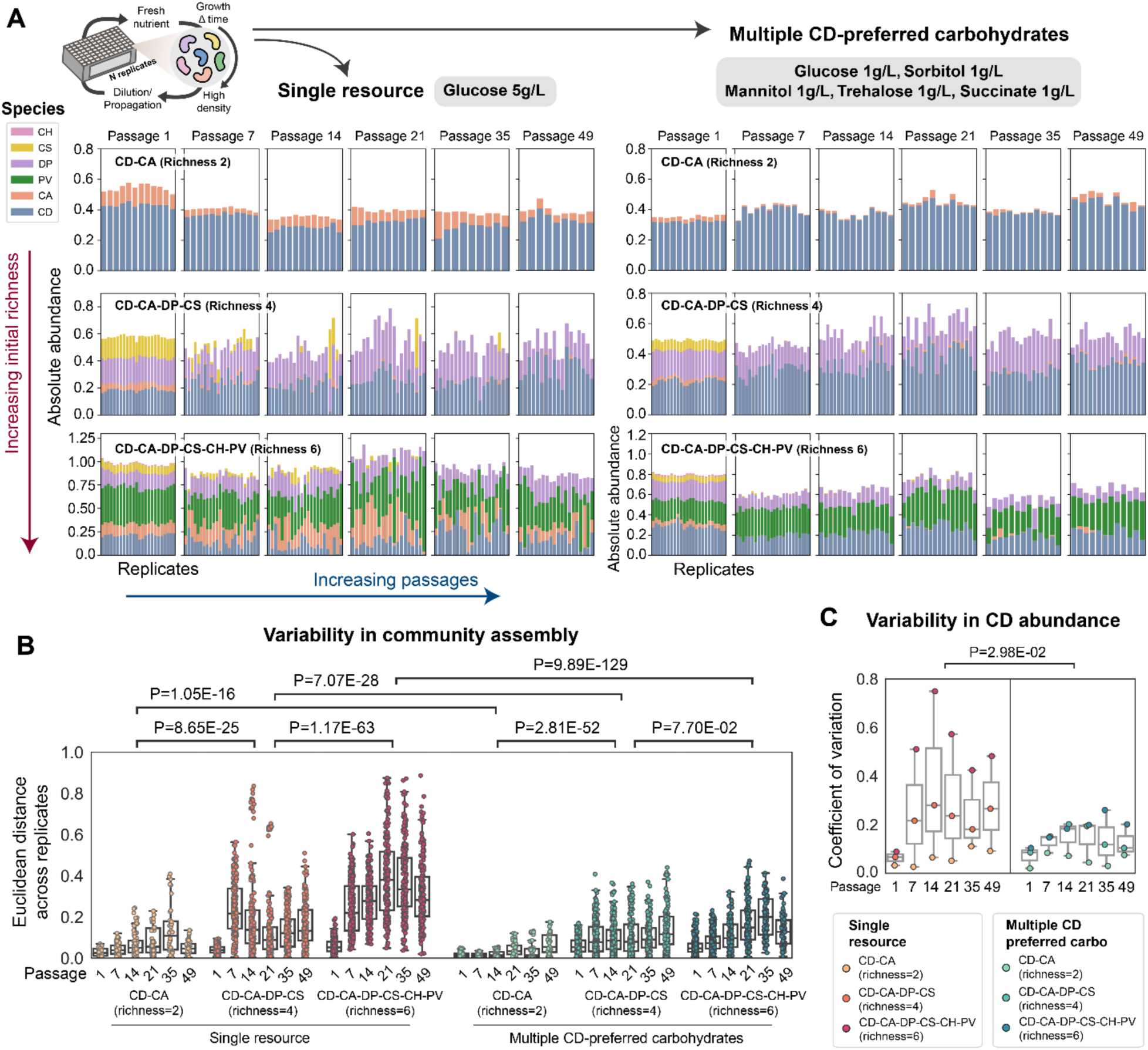
Nutrient environments shape the variability of long-term community assembly. **a**, Stacked bar plot of the absolute abundance (OD600) of the independent replicates of 2, 4, and 6-member community containing *C. difficile* over 49 passages grown in media containing only glucose (**left**) or multiple *C. difficile*-preferred carbohydrates (**right**). Two-member communities were grown with 12 independent replicates, whereas 4 and 6-member communities were grown with 24 independent replicates. Contaminated replicates were excluded from the analysis. **b**, Euclidean distance of community composition between pairs of replicates from the 1^st^ to the 49^th^ passages in the two media conditions. *p*-values from unpaired *t*-test (two-sided) between different sample groups are shown. **c**, Coefficient of variations of *C. difficile* abundance across replicates from the 1^st^ to the 49^th^ passages in the two media conditions. The *p*-value from unpaired *t*-test (two-sided) of the coefficient of variations between the two media conditions is shown.

To evaluate whether species richness in a community also influences the degree of variability, we randomly assembled communities with different species richness. The Euclidean distances of species relative abundance between all pairs of replicates increased over time more consistently in the media with multiple *C. difficile*-preferred carbohydrates. In media containing only glucose, the Euclidean distance displayed non-monotonic trends over time (**Fig. 5b**). Overall, the Euclidean distances were higher in communities with higher species richness in both media types, and in the media containing only glucose than in media with multiple *C. difficile*-preferred carbohydrates. Although *C. difficile* abundance was lower in the glucose-only media ^55^, the variability in its abundance was higher than in multiple preferred carbohydrates (**Fig. 5c, S19b**). This highlights a potential trade-off between high variability versus strong inhibition of *C. difficile* due to variations in the nutrient landscape.

Variability in propagule pressure of *C. difficile* (i.e. initial abundance in the community at the beginning of each passage) could contribute to the variability in abundance across repetitive regrowth cycles ^14^. To determine how nutrient environment shapes the sensitivity to variation in propagule pressure, we characterized community assembly in the glucose-only media versus multiple *C. difficile-*preferred carbohydrates media (**Fig. S19c**). In the glucose-only media, *C. difficile* growth displays a larger change as a function of its initial amount in low to medium-richness communities compared to the multiple *C. difficile*-preferred carbohydrates media, i.e. more sensitive to propagule pressure (**Fig. S19d**). Thus, the sensitivity to initial abundance in different nutrient environments can provide insights into the observed variability in community dynamics.

## DISCUSSION

The community-acquired incidence of CDI continues to rise despite the expanding treatment options including FMT and emerging defined bacterial therapeutics ^99–101^. *C. difficile* can colonize individuals for yearlong timescales ^6,8,10,102^ and act as a reservoir for *C. difficile,* increasing the chance of infection in the host ^16^ and other individuals via transmission ^8,19–21^. Identifying strategies that reduce *C. difficile*’s ability to persist in the human gut could potentially reduce the prevalence and severity of CDI. Human gut microbiota interactions are major variables influencing colonization ability. However, we lack an understanding of the molecular and ecological mechanisms shaping *C. difficile* growth in communities over long timescales. We exploited high-throughput *in vitro* experiments combined with dynamic ecological modeling to understand the role of community context on the ability of *C. difficile* to coexist with diverse human gut bacteria and elucidate factors shaping variability in its growth over hundreds of generations. We identified key species that display stable coexistence or unstable dynamics with *C. difficile*, consistent with the quantitative effects of their inferred inter-species interactions on community dynamics. Our findings yield insights into the ability of *C. difficile* to persist in different gut communities, which could aid in devising strategies to reduce its persistence. Importantly, we found that prolonged co-culture of *C. difficile* with PV could lead to *C. difficile* evolutionary adaptation that reduces its virulence through metabolic alterations. This opens up a new avenue for *C. difficile* interventions where bacterial therapeutics or diets could be designed to steer *C. difficile* evolutionary adaptations towards reduced colonization ability and attenuated virulence.

The variability of *C. difficile* across individuals could influence transmission and disease outcomes. For example, higher variability generates higher uncertainty of *C. difficile* growth or toxin production and could reduce the predictability and efficacy of treatments for CDI. We demonstrated that certain pairs of species have more unstable dynamics across different community contexts and nutrient environments (**Fig. 1f, S5h-j**), which leads to higher variability in growth. Further, nutrient environments and the number of species in the community can also influence variability in *C. difficile* growth over long timescales. Using a library of isolates from soil samples and *Caenorhabditis elegans* intestine, a previous study ^103^ demonstrated that increasing the number of species in the community or nutrient concentrations ^104,105^ reduced the stability of community dynamics, shifting the system from stable coexistence to persistent fluctuations. In this study, we observed that communities with higher species richness displayed higher variability in abundances over long timescales (**Fig. 5**). While fixing the total amount of carbon, there was higher variability in community composition and *C. difficile* abundance in an environment containing a single accessible resource as opposed to multiple distinct resources. An environment with a single accessible resource may enhance the strength of resource competition and/or production of toxic metabolic byproducts. While *C. difficile*’s abundance was reduced in this environment, its variability was enhanced, highlighting a potential trade-off.

The human gut microbiome is dominated by *Bacteroides* species such as PV ^75–78^ that stably engraft for long timescales ^106–108^. PV has been highlighted as a potential candidate for live bacterial therapeutics to treat recurrent CDI (rCDI) due to its high engraftment efficiency ^108^. Supporting this notion, PV was a dominant member in a 7-member bacterial community in germ-free mice (∼50% fraction across all conditions) and maintained a high relative abundance for up to 2 weeks even after the mice were challenged with *C. difficile* (**Fig. S17**). Understanding how highly abundant and stable members of the human gut microbiome coexist and interact with *C. difficile* over the long term could have important therapeutic implications. Our results demonstrated that decreasing the concentration of carbohydrates shifted PV and *C. difficile* community dynamics from competitive exclusion to coexistence over 341 generations (**Fig. 2, S5e-f**). Using exo-metabolomic profiling, we revealed that this coexistence can be explained by a high degree of metabolite cross-feeding from PV to *C. difficile*, including major amino acids that fuel *C. difficile*’s Stickland metabolism. This cross-feeding of amino acids from *Bacteroides* to *Clostridium* species has been previously reported ^82^, and there are other *in vitro* and *in silico* evidence of amino acids cross-feeding among gut microbes ^82,109^. Our results suggest that cross-feeding was not present in the presence of high carbohydrate concentrations, consistent with a previous study showing that high concentrations of acetate suppress the release of amino acids by *Bacteroides* species ^83^. Notably, a moderate reduction in the concentrations of key resources could massively alter the interactions between PV and *C. difficile*. This implies that diet could be precisely manipulated to alter interactions between gut microbes and *C. difficile* and influence the evolutionary trajectories of *C. difficile*.

Persistent colonization enables *C. difficile* to adapt to changing environments within the human gut. Evolutionary adaptations of *C. difficile* could impact the host through alterations in metabolism and virulence ^29–32^. Our current knowledge of *C. difficile*’s evolution is largely based on retrospective sequencing-based analyses ^110^ and genome comparisons to reveal genetic changes predicted to alter key phenotypes ^29,31,32,111–113^. We lack an understanding of the role of human gut microbiota inter-species interactions on *C. difficile* evolutionary adaptations. Our study demonstrated that prolonged coexistence between *C. difficile* and PV yielded evolutionary adaptations in *C. difficile* that shifted its metabolism from consuming proline to glucose and altered its fitness under different concentrations of resources (**Fig. 3, S13-14, Table S4**). These metabolic changes enhanced its ability to coexist with PV (**Fig. S9b**). Although a previous study showed that *C. difficile* strains with large genotypic variations have minimal differences in inter-species interactions with human gut bacteria ^55^, here we demonstrate that two single-point mutations in *C. difficile* metabolic genes could substantially alter the interaction networks with human gut microbiota (**Fig. 4a-d**). This implies that although human gut microbiota inter-species interactions with *C. difficile* are generally robust to genotypic variations, a small number of mutations in key genes could yield large changes in the interaction network.

An important question is whether this metabolic adaptation of *C. difficile* alters disease severity or propensity for CDI. Proline metabolism of *C. difficile* has been shown to affect toxin production ^79^ and colonization of the murine gut ^80^. Indeed, our evolved *C. difficile* strain displayed a substantial reduction in toxin production and colonization ability, thus ameliorating disease severity in the murine gut compared to the ancestral *C. difficile* strain (**Fig. 4e-i**). Deletion of *prdB* in *C. difficile,* an essential enzyme in the proline Stickland fermentation pathway ^79^, reduced colonization and toxin production in the murine gut ^80^. This is consistent with the reduced disease severity observed in mice colonized with the evolved *C. difficile* strain which has evolved to use less proline. These results demonstrate that proline utilization plays a critical role in the colonization and infection process. Our study highlights how two-point mutations in the *C. difficile* genome from evolutionary adaptation can substantially alter virulence and colonization ability. This implies that *C. difficile*’s pathogenic potential displays fragility to mutations in specific genes. *C. difficile* clinical isolates have displayed variable effects on mice ^114^ and avirulent strains have been shown to protect against CDI by outcompeting the virulent *C. difficile* strains ^115^. As opposed to eliminating *C. difficile* from the gut, future studies could investigate interventions that steer *C. difficile* metabolic states and suppress its virulence (e.g. shifting away from proline utilization) while also promoting colonization and fitness in the gut, thus protecting against virulent *C. difficile* strains.

## Materials and Methods

### Strain, media, and growth conditions

The strains used in this work were obtained from the sources listed in **Table S1**. The non-toxigenic *C. difficile* isolate MS001 was obtained from a previous study ^55^. Single-use glycerol stocks were prepared as described previously ^51^. The media used in this work are anaerobic basal broth (ABB, Oxoid) for growing starter cultures, and in-house Defined Media (DM) formulated based on previous study ^51^ (recipe in **Table S2**).

For all experiments, cells were cultured in an anaerobic chamber (Coy Lab products) with an atmosphere of 2.0 ± 0.5% H2, 15 ± 1% CO2, and balance N2 at 37 C. Starter cultures were inoculated by adding 200 μl of a single-use 25% glycerol stock to 5 ml of anaerobic basal broth media (ABB) and grown at 37 C without shaking.

### Long-term growth experiments

Starter cultures of *C. difficile* and gut commensal bacteria were prepared. For experiments in **Fig. 1**, the media used are DM, DM with 75% less carbohydrate concentration, and DM with 100% higher amino acid concentration. For experiments in **Fig. 5**, the media used are DM containing only 5 g/L glucose as a sole carbohydrate source, and DM containing 1 g/L glucose, 1 g/L sorbitol, 1 g/L mannitol, 1 g/L trehalose, and 1 g/L succinate as carbohydrate sources.

The cell pellets from starter cultures were collected by centrifugation at 3,000 x g for 5 min, and then washed with experimental media. The washed cell pellets were resuspended into the experimental media to a final OD600 of approximately 0.1. To assemble communities, the monocultures of *C. difficile* and each gut species were mixed in equal proportions based on OD600 and inoculated into a 2 mL 96-deep-well plate (Nest Scientific) containing experimental media to an initial OD600 of 0.01. For instance, the OD600 of each species in two-member and 6-member communities are 0.01/2=0.005 and 0.01/6=0.00167 respectively. These plates were covered with gas permeable seal (Breathe-Easy^®^ sealing membrane) and incubated at 37 °C anaerobically. After every 24 hours, OD600 was measured with a Tecan F200, and the cells were passaged with 40X dilution to new 96-deep-well plates containing experimental media. The monocultures of individual species in the community were also inoculated and passaged as a control and to monitor for changes in carrying capacity. Every 7 days, aliquots of the cultures were preserved as glycerol stocks and cell pellets were collected for DNA extraction, PCR amplification, and NGS sequencing. Depending on the experiment, the communities were maintained for 35 to 64 passages. We calculated the number of generations per passage as the log2 of the dilution factor ^116^. Thus, we estimate that the communities were maintained for up to 341 generations, which is long enough for adaptation to occur based on previous studies ^41,42^.

### Isolation of *C. difficile* strains from communities

Communities from the evolution experiments were preserved as glycerol stocks. Aliquots from the glycerol stock were streaked into *C. difficile* selective plates to isolate single colonies, the plates were incubated at 37°C for 24 h, and the *C. difficile* strain was grown in a liquid culture from a single colony.

*C. difficile* selective plates were prepared by autoclaving *C. difficile* agar (Oxoid CM0601) and adding horse blood (Lampire 7233401, 70 mL/1L media), norfloxacin (Santa Cruz 215586, 120 μg/mL), moxalactam (Santa Cruz 250419, 320 μg/mL), and erythromycin (Santa Cruz 204742, 100 μg/mL) after the media cooled to 55°C.

### Fluorescence microscopy of *C. difficile*

Starter cultures of the evolved and ancestral *C. difficile* strain were prepared. The cell pellets from starter cultures were collected by centrifugation at 3,000 x g for 10 min, and then washed with DM. The washed cell pellets were resuspended into DM to a final OD600 of approximately 0.1. These cultures were inoculated into new culture tubes containing DM to an initial OD600 of 0.01 by adding 500 μl of washed starter cultures to 4.5 mL media. After 6 h and 24 h of growth, 100 μl aliquots were taken, stained with SYBR Green dye, and viewed with a microscope (Nikon Eclipse Ti-E inverted microscope) at 20× dry objective with appropriate filter sets. Images were captured with Photometrics CoolSNAP Dyno CCD camera and associated software (NIS-Elements Ver. 4.51.00).

### Logistic growth model

The logistic growth model was used to describe population growth dynamics in monoculture experiments. The logistic growth model for species *i* takes the following form:

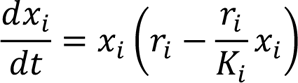

where *x_i_* is the absolute abundance of species *i*, parameter *r_i_* is its maximum growth rate, and *K_i_* is its carrying capacity. We cut time points where the OD600 drops below > 10% to exclude the death phase. Thus, the steady-state solution of the model is the carrying capacity (*K* ) (i.e. the value of *x_i_* when 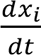 equals 0). We also excluded data points less than 120% of the initial OD600 (OD600 at t=0) to exclude the lag phase which is not captured in the logistic model. A custom MATLAB script is used to estimate the parameters θ_*i*_ = [*r_i_*, *K_i_*] in the logistic growth model. For each species *i*, the model is fitted to experimental data with L2 regularization. Specifically, given a series of *m* experimental OD600 measurements, 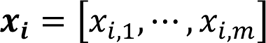, and a series of OD600 simulated using parameter θ_*i*_ at the same time intervals, 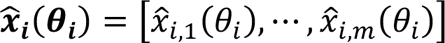, the optimization scheme minimizes the cost function:

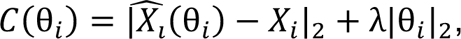

where λ is the L2 regularization parameter and | ⋅ |_2_ indicates vector 2-norm. To find a suitable regularization parameter λ, we took λ values from the set (10^-3^, 3 × 10^-3^, 10^-2^, 3 × 10^-2^, 0.1, 0.3, 1, 3, 10). Based on the cost for each species as a function of λ, we picked regularization parameter λ = 0.03. Solutions to the logistic growth model were obtained using the ode15s solver and the optimization problem was solved using FMINCON in MATLAB (R2022a).

### Bacterial genome DNA extraction and next-generation sequencing

All the genomic DNA (gDNA) extraction and next-generation sequencing sample preparation were performed as described previously ^51,55^. Bacterial gDNA extractions were carried out using a modified version of the Qiagen DNeasy Blood and Tissue Kit protocol in 96-well plates. Briefly, cell pellets were resuspended in 180-μl enzymatic lysis buffer containing 20 mg/ml lysozyme (Sigma-Aldrich), 20 mM Tris–HCl pH 8 (Invitrogen), 2 mM EDTA (Sigma-Aldrich), and 1.2% Triton X-100 (Sigma-Aldrich), and then incubated at 37°C at 600 RPM for 30 min. Samples were treated with 25 μL 20 mg/ml Proteinase K (VWR) and 200 μL buffer AL (Qiagen), mixed by pipette, and then incubated at 56°C at 600 RPM for 30 min. Samples were treated with 200 μL 200 proof ethanol (Koptec), mixed by pipette, and transferred to 96-well nucleic acid binding plates (Pall). After washing with 500 μL buffer AW1 and AW2 (Qiagen), a vacuum was applied for 10 min to dry excess ethanol. Genomic DNA was eluted with 110 μL buffer AE (Qiagen) preheated to 56°C and then stored at −20°C.

Genomic DNA concentrations were measured using the Quant-iT™ dsDNA Assay Kit (Invitrogen) with a 6-point DNA standard curve (0, 0.5, 1, 2, 4, 6 ng/μL biotium). 1 μL of samples and 5 μL of standards were diluted into 95 μL of 1× SYBR green (Invitrogen) in TE buffer and mixed by pipette. Fluorescence was measured with an excitation/emission of 485/535 nm (Tecan Spark). Genomic DNA was then normalized to 2 ng/µL by diluting in molecular grade water (VWR International) using a Tecan Evo Liquid Handling Robot.

Dual-indexed primers for multiplexed amplicon sequencing of the V3-V4 region of the 16S rRNA gene were designed as described previously ^51,60^. PCR was performed using the normalized gDNA as template and Phusion High-Fidelity DNA Polymerase (Thermo Fisher) for 25 cycles with 0.05 μM of each primer. Samples were pooled by plate, purified using the DNA Clean & Concentrator™-5 kit (Zymo) and eluted in water, quantified by NanoDrop, and combined in equal proportions into a library. The library was quantified using Qubit 1× HS Assay (Invitrogen), diluted to 4.2 nM, and loaded at 10 pM onto Illumina MiSeq platform for 300-bp paired-end sequencing using MiSeq Reagent Kit v2 (500-cycle), or loaded at 21 pM using MiSeq Reagent Kit v3 (600-cycle) depending on the desired sequencing reads.

### Next-generation sequencing data analysis to determine community composition

Sequencing data were analyzed as described previously ^55,60^. Briefly, reads were demultiplexed with Basespace FastQ Generation, and the FastQ files were analyzed using custom Python scripts. Paired reads were merged using PEAR (Paired-End reAd mergeR) v0.9.0 ^117^. A reference database containing 16S V3-V4 region of each species in the study was created by assembling consensus sequence based on sequencing results of each monospecies. Reads were mapped to the reference database using the mothur v1.40.5 command classify.seqs using the Wang method with bootstrap cutoff value of 60% ^118,119^. Relative abundance was calculated by dividing the read counts mapped to each organism by the total reads in the sample. Absolute abundance was calculated by multiplying the relative abundance of an organism by the OD600 of the sample as previously described ^14,60^. Samples were excluded from further analysis if > 1% of the reads were assigned to a species not expected to be in the community (indicating contamination).

### generalized Lotka-Volterra models for fitting non-passaging data

The generalized Lotka-Volterra (gLV) model is a set of coupled ordinary differential equations that describe the growth of interacting species over time,

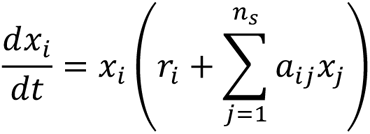

where *x_i_*is the abundance of species *i* and *n*_2_is the total number of species. Model parameters that need to be estimated from data include the species growth rate, denoted as *r_i_* , and coefficients that determine how species *j* affects the growth of species *i* , denoted as *a*_!j_. The data used for parameter estimation is the growth of species over time under different inoculation conditions. For monoculture growth data, we use OD600 measurements only, whereas for community data, this was obtained by multiplying the relative abundance obtained from 16S sequencing by the total OD600. The dataset used to fit the gLV model is DATASET002 (**Table S3**).

A prior over the parameter distribution is set so that growth rates have a mean of 0.3, self-interaction terms have a mean of -1, and inter-species interaction terms have a mean of -0.1. Given a dataset of measured species abundances over time after inoculating different combinations of species, the model parameters are determined by minimizing a cost function given by a weighted squared difference between model-predicted species abundances and measured abundances and a penalty for deviations from the prior mean. Using the fitted parameter estimates, the covariance of the posterior parameter distribution is approximated as the inverse of the Hessian (matrix of second derivatives) of the cost function with respect to the model parameters. The Expectation-Maximization (EM) algorithm is used to optimize the precision of the prior parameter distribution and the precision of the noise distribution, which collectively determine the degree to which estimated parameters are penalized for deviations from the prior mean^120^. In other words, the precision of the prior and noise are hyperparameters that determine the degree of regularization. To evaluate model prediction performance on held-out data, we performed 10-fold cross validation where the degree of regularization was optimized using the EM algorithm and only community samples were subjected to testing (i.e. monoculture data was reserved only for model training). See **Supplementary Text** for a more detailed description of parameter estimation and the EM algorithm.

### Generalized Lotka-Volterra models for fitting long-term passaging data

A general Lotka-Volterra (gLV) model with elastic net regularization was used to simulate long-term growth experiments with passaging (repetitive regrowth cycles):

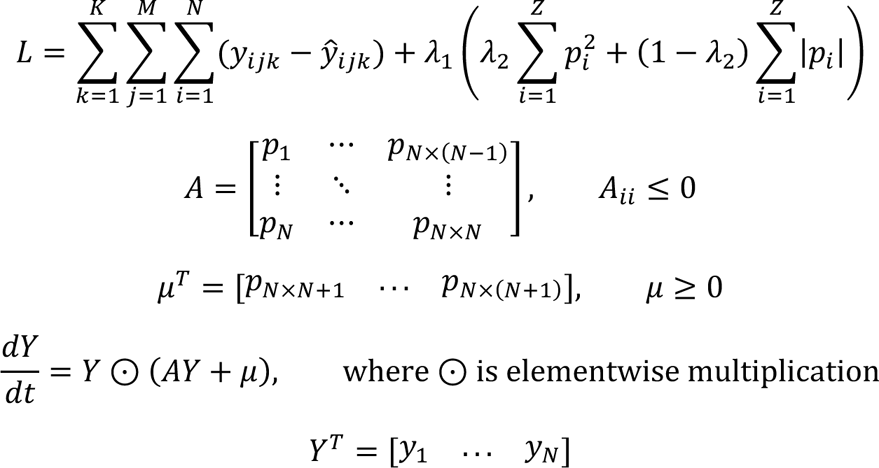

where ŷ is the predicted output, *y* is the observed output or species normalized optical density (OD), *K* is the number of passages, *M* is the number of averaged experimental samples, *N* is the number of species, *λ*_1_ = 10^-6^ is the overall elastic net regularization parameter, *λ*_2_ = 0.5 is the parameter weighing between Ridge and Lasso regularization, *p_i_* are the general Lotka-Volterra (gLV) parameters, *A* is the inferred gLV interaction matrix, and *μ* are the inferred gLV growth rates. To obtain the predictions, we integrated *Y* over a time span [0, *t_f_*], where *t_f_* = 24, using a fixed-step Euler’s method for simplicity and applied a 40-fold dilution to the end time point of each integration *T* times for *T* passages. The loss was optimized using a global constrained optimization algorithm (Low Discrepancy Sequence Multi-Level Single-Linkage) with a population of 32 samples and a local optimizer Sbplx, a more robust Nelder-Mead method ^121–124^. The diagonals of matrix *A* are constrained to be negative, and *μ* was constrained to be nonnegative. We fitted 64 models over the entire experimental dataset, and unstable models were discarded to obtain final parameter estimates, for which distribution is assumed Gaussian. The gradient-free optimization method and Euler’s method were chosen in favor of gradient-based optimization and adaptive higher-order ordinary differential equation solvers to handle gLV model instability due to long integration times. The dataset used to fit the gLV model is DATASET001 (**Table S3**).

### Whole-genome sequencing and variant calling

We subjected six strains for whole-genome sequencing: Ancestral *C. difficile* DSM 27147 and ancestral *P. vulgatus* ATCC 8482 strain, *C. difficile* and PV strain that were passaged alone in the media with reduced carbohydrates concentration*, a*nd *C. difficile* and PV strain isolated from CD-PV co-culture after 64 passages in the media with reduced carbohydrates concentration. Cultures were streaked into *C. difficile* selective agar plate (for isolating *C. difficile*) or ABB agar plate (for isolating PV) to isolate single colonies. Although ABB is not selective for PV, its proportion is much higher than *C. difficile* in the CD-PV co-culture and thus has a much higher number of colonies in the ABB agar plate. For all conditions, one colony was isolated, grown to OD600 of 0.3, and subjected to whole-genome sequencing. Besides the six strains, we also subjected the whole CD-PV population (all 3 biological replicates) after 35, 49, and 64 passages in the media with reduced carbohydrate concentration to whole-genome sequencing.

The cultures were centrifuged to obtain the cell pellets. Genomic DNA was extracted using Qiagen DNeasy Blood and Tissue Kit according to the manufacturer’s protocol. The harvested DNA was detected by the agarose gel electrophoresis and quantified by a Qubit fluorometer. The genomic DNA was sent to SeqCenter (Pittsburgh, PA, USA) for paired-ends Illumina sequencing. Sample libraries were prepared using the Illumina DNA Prep kit and IDT 10 bp UDI indices, and sequenced on an Illumina NextSeq 2000, producing 2 x 151 bp reads. Demultiplexing, quality control, and adapter trimming were performed with bcl-convert (v3.9.3) Illumina software.

To identify mutations in the six strains, we performed variant calling analysis using BreSeq version 0.38.1 ^125^ with default settings using *C. difficile* R20291 reference genome (GenBank ID: FN545816.1) or *P. vulgatus* ATCC 8482 reference genome, and performed a genomic comparison between the ancestral strain and strains after passaging both in monoculture and co-culture conditions. We only detected two non-synonymous single-point mutations in *gene 206* (G533W) and in *prdR* (A341V) for *C. difficile* from the CD-PV pair, and no mutations in *C. difficile* that was passaged alone, PV that was passaged alone, or PV from CD-PV pair. To identify lower abundance mutations in the CD-PV population, we performed variant calling analysis using Snippy V4.6.0 ^126^. To estimate the proportion of mutants in the population, we used the ratio of the number of alternate reads (reads of the mutation) to the total number of reads at the locus (number of alternate reads + number of reference reads) extracted from Snippy vcf result files.

### Sanger sequencing of the *prdR* gene

To check the presence of the *prdR* A341V mutation in other biological replicates of CD-PV co-culture and other pairwise communities, we performed Sanger sequencing on *C. difficile* isolated from two other biological replicates of the CD-PV pair after 64 passages, three biological replicates of the CD-BU pair after 64 passages, two biological replicates of the CD-CA pair after 42 passages, one biological replicate of the CD-DP pair after 49 passages, one biological replicate of the CD-DP pair after 64 passages, and two biological replicates of the CD-EL pair after 64 passages (one colony each).

Aliquots from the glycerol stock of communities at the end of the passaging experiments were streaked into *C. difficile* selective plates to isolate single colonies. The *C. difficile* strains were grown from a single colony to OD600 of 0.3. The cultures were centrifuged to obtain the cell pellets. Genomic DNA was extracted using Qiagen DNeasy Blood and Tissue Kit according to the manufacturer’s protocol.

PCR was performed using the gDNA as a template (2 ng/µL) and Phusion High-Fidelity DNA Polymerase (Thermo Fisher) for 25 cycles with 0.2 μM of each primer. The primers were designed to amplify 526 bp of the *prdR* gene targeting the mutated region (A341V). The primer sequence is *F*: CAGAAGCTAAGATATTAGCTCTTGAA and *R*: ATTGGTAGCTGATATTATTCTAGGA. The amplified PCR product was subjected to Sanger Sequencing (Functional Biosciences).

### Transcriptomic profiling

Ancestral *C. difficile* monoculture, evolved *C. difficile* monoculture, CD ancestral + PV coculture, and CD evolved + PV coculture conditions were inoculated from starter cultures into individual culture tubes containing DM. For monoculture conditions, *C. difficile* was inoculated to an OD600 of 0.01. For cocultures, *C. difficile* and PV were inoculated to an equal ratio (OD600 of 0.005 each). The cultures were incubated anaerobically at 37°C with no shaking for ∼6 h until the culture reached the exponential phase (OD600 ∼0.2). 1000 μL of the culture was taken for OD600 measurement and total DNA extraction for next-generation sequencing, and 2000 μL of the culture was taken for total RNA extraction for transcriptomics. 4000 μL of RNAprotect (Qiagen) was added to 2000 μL of culture and incubated for 5 min at room temperature. Cultures were then centrifuged at room temperature for 10 min at 3000 g and the supernatant was carefully removed. Cell pellets were immediately subjected to RNA extraction using acidic phenol bead-beating method. Pellets were resuspended in 500 μL 2× Buffer B (200 mM sodium chloride, 20 mM ethylenediaminetetraacetic acid) and transferred to 2 mL microcentrifuge tubes containing 500 μL Phenol:Chloroform:IAA (125:24:1, pH 4.5) and 210 μL 20% sodium dodecyl sulfate and were bead-beated with acid washed beads (Sigma G1277) for 3 min. All solutions used for RNA extraction were RNAse-free. Samples were centrifuged at 4°C for 5 min at 7,200 g, and 600 μL of the upper aqueous phase was added to 60 μL 3 M sodium acetate and 660 μL cold isopropanol and chilled on ice for 5 min before freezing for 5 min at −80°C. Samples were centrifuged at 4°C for 15 min at 18,200 g, the supernatant was decanted, and the pellet was washed with cold 100% ethanol. The pellets were dried in a biosafety cabinet for 15 min and then resuspended in 100 μL RNAse-free water. Samples were purified using RNeasy Mini Kit (Qiagen) and genomic DNA was removed using RNAse-Free DNase Set (Qiagen). Two replicates of each condition were sent to Novogene Corporation Inc (Sacramaneto, CA, United States of America) for rRNA depletion, cDNA library preparation, and sequencing on Illumina NovaSeq. Data was de-multiplexed using Illumina’s bcl2fastq 2.17 software, where one mismatch was allowed for index sequence identification.

The compressed FASTQ files were quality-checked using the FastQC tool v0.12.1^127^. The BBDuk, BBSplit, and BBMap tools from BBTools suite (v38.42) ^128^ were used to trim adapters, deplete rRNA, and map the remaining mRNA reads to the reference genomes. For monoculture or cocultures containing *C. difficile*, the reference genome was obtained from GenBank (FN545816.1). The feature-Counts package v1.6.4 ^129^ from the SubRead suite was used to map reads to features on the genome and quantify raw counts for each transcript. Reads per kilobase million (RPKM) values were computed using a custom Python script to see the agreement of gene expression between biological replicates. The gene expression (represented by RPKM values) shows a good correlation between biological replicates (Pearson’s R=0.97-0.98, P<10E-05) (**Fig. S11b**). The DESeq2 Bioconductor library v4.0.3 ^130^ was used in R v4.0.4 to quantify differential gene expression using a negative binomial generalized linear models with apeglm shrinkage estimator ^131^. When calculating RPKM of *C. difficile* genes in the CD-PV cocultures, the “reads mapped” in the denominator was the number of reads mapped to the *C. difficile* genome. Similarly, when quantifying differential gene expression for *C. difficile* genes in the CD-PV cocultures, only reads mapped to the *C. difficile* genome were provided to DeSeq2. We define differentially expressed genes (DEGs) as those with >2-fold change and a *p*-value less than 0.05.

### *C. difficile* toxin measurements using ELISA

Toxin (both TcdA and TcdB) concentrations were determined in the ancestral and evolved *C. difficile* strains by comparison to a standard curve using ELISA (tgcBiomics, Germany). The blank media used to grow the cultures were also included in the assay to measure any background noise. All the samples subjected to toxin measurements in this study were processed in parallel at the same time using the same batch of ELISA kits to minimize batch-to-batch variations and ensure comparable results.

### Exo-metabolomic profiling

Starter cultures of *C. difficile* and gut bacteria were prepared. The cell pellets from starter cultures were collected by centrifugation at 3,000 x g for 5 min, and then washed with either DM or DM with reduced carbohydrate concentration. The washed cell pellets were resuspended into either DM or DM with reduced carbohydrate concentration to a final OD600 of approximately 0.1. For monocultures, bacteria were inoculated to an OD600 of 0.01 in media containing either DM or DM with reduced carbohydrate concentration. For co-cultures, *C. difficile* and PV were inoculated to an equal ratio (OD600 of 0.005 each) in media containing either DM or DM with reduced carbohydrate concentration. The cultures were incubated at 37°C anaerobically. Three biological replicates were performed for each sample. At specific time points (6, 12, or 24 hours), the cultures were centrifuged at 3,000 x g for 10 min, and the supernatants were filter sterilized.

For metabolite extraction, 25 µL of each sterilized supernatants sample were pipetted into a microcentrifuge tube. The extraction solvent consisted of 1:1 methanol:ethanol containing 22 µM D4-succinate (Sigma 293075). 112.5 µL of the extraction solvent was added to each sample, followed by a 10 min incubation on ice. Then, 87.5 µL of molecular biology-grade water was added to the extraction tube, followed by another 10 min incubation on ice. The samples were centrifuged at 16,000 x g for 10 min at 4°C, and the supernatant (190 µL) was transferred to a Captiva cartridge (Agilent 5190-1002, 1mL). The sample was allowed to flow through under vacuum, followed by two elutions with 250 µL of 2:1:1 water:methanol:ethanol. Both the flowthrough and the elutions were collected in one tube and evaporated under vacuum at 45°C for 2 h. The dried metabolite pellet was stored at -80°C until it was resuspended in 70:20:10 acetonitrile:water:methanol (100 µL) prior to LC/MS analysis.

Extracts were separated on an Agilent 1290 Infinity II Bio LC System using an InfinityLab Poroshell 120 HILIC-Z column (Agilent 683775-924, 2.7 µm, 2.1 x 150 mm), maintained at 15°C. The chromatography gradient included mobile phase A containing 20 mM ammonium acetate in water (pH 9.3) and 5 µM of medronic acid, and mobile phase B containing acetonitrile. The mobile phase gradient started with 10% mobile phase A increased to 22% over 8 min, increased to 40% by 12 min, 90% by 15 min, and then held at 90% until 18 min before re-equilibration at 10% (held until 23 min). The flow rate was maintained at 0.4 mL/min for most of the run, but increased to 0.5 mL/min from 19.1 min to 22.1 min. The UHPLC system was connected to an Agilent 6595C QqQ MS dual AJS ESI mass spectrometer. This method was operated in polarity-switching mode. The gas temperature was kept at 200°C with flow at 14 L/min. The nebulizer was at 50 PSI, sheath gas temperature at 375°C, and sheath gas flow at 12 L/min. The VCap voltage was set at 3000V, iFunnel high pressure RF was set to 150 V, and iFunnel low pressure RF was set to 60 V in positive mode. In negative mode, the VCap voltage was set to 2500 V, the iFunnel high pressure RF was set to 60 V, and iFunnel low pressure RF was set to 60 V. A dMRM inclusion list was used to individually optimize fragmentation parameters. The injection volume was 1 µL.

Raw data was collected in .d format and checked manually in Agilent MassHunter Qualitative Analysis. The data was then uploaded to Agilent MassHunter Quantitative Analysis for quantitation using relative internal standard calculations to calculate analyte concentrations. After manual inspection and integration, analyte concentration (ng/mL of reconstituted extract) was exported to .csv files.

### HPLC quantification of organic acids

Starter cultures of ancestral *C. difficile*, evolved *C. difficile*, and PV were prepared. The cell pellets from starter cultures were collected by centrifugation at 3,000 x g for 5 min, and then washed with DM or DM with reduced concentration of carbohydrates. The washed cell pellets were resuspended into the respective media to a final OD600 of approximately 0.1, inoculated into a 5 mL culture tubes to an initial OD600 of 0.01, and incubated at 37 °C anaerobically. For co-cultures, *C. difficile* and PV were inoculated to an equal ratio (OD600 of 0.005 each). Three biological replicates were performed for each sample.

After 24 hours, the cultures were centrifuged at 3,000 x g for 10 min, and the supernatants were filter sterilized. Then, 2 μL of H2SO4 was added to the supernatant samples to precipitate any components that might be incompatible with the running buffer. The samples were then centrifuged at 3,000 x g for 10 min and then 150 μL of each sample was filtered through a 0.2 μm filter using a vacuum manifold before transferring 70 μL of each sample to an HPLC vial. HPLC analysis was performed using a Shimadzu HPLC system equipped with a SPD-20AV UV detector (210 nm). Compounds were separated on a 250 × 4.6 mm Rezex^©^ ROA-Organic acid LC column (Phenomenex Torrance, CA) run with a flow rate of 0.2 mL min^−1^ and at a column temperature of 50 °C. The samples were held at 4 °C prior to injection. Separation was isocratic with a mobile phase of HPLC grade water acidified with 0.015 N H2SO4 (415 µL L^−1^). At least two standard sets were run along with each sample set. Standards were 100, 20, 4, and 0.8 mM concentrations of butyrate, succinate, lactate, and acetate, respectively. The injection volume for both sample and standard was 25 µL. The resultant data was analyzed using the Shimadzu LabSolutions software package.

### Hill function fitting

The sensitivity of *C. difficile* growth to proline or glucose concentration was quantified by fitting the data to the Hill equation:

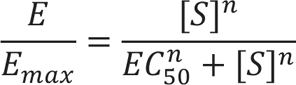

where *E* is the normalized Area Under the Curve of *C. difficile* growth for 24 h (AUC24h), *E_max_* is the maximum normalized AUC24h across all proline/glucose concentrations, [*S*] is the proline/glucose concentration, *EC*_50_ is the proline/glucose concentration that produces 50% of *E_max_* value, and *n* is a measure of ultrasensitivity. The data were fit using the curve_fit function of the scipy package optimization module in Python.

### Gnotobiotic mouse experiments

All germ-free mouse experiments were performed following protocols approved by the University of Wisconsin-Madison Animal Care and Use Committee. We used 12-week-old C57BL/6 gnotobiotic male mice (wild-type) and a regular diet (Chow diet, Purina, LabDiet 5021). All bacterial strains were grown at 37 °C anaerobically in Anaerobe Basal Broth (ABB, Oxoid) to stationary phase. Commensal gut bacteria strains for oral gavage were mixed in equal proportions based on OD600, whereas *C. difficile* strains for oral gavage were diluted to an OD600 corresponding to ∼50,000 CFU/mL based on prior conversion calculation from CFU counting. These cultures were transferred to Hungate tubes (Chemglass) on ice prior to oral gavage. To verify and ensure similar dosage for ancestral *C. difficile* and evolved *C. difficile* gavage, aliquots of *C. difficile* cultures were plated on agar plates to calculate the exact CFU. The CFU for ancestral *C. difficile* strain was 50,400 CFU/mL, whereas the CFU for evolved *C. difficile* strain was 49,200 CFU/mL. On day 0, 0.2 mL of commensal gut bacteria (core community) was introduced into the mice by oral gavage inside a Biological Safety Cabinet (BSC) and the mice were housed in biocontainment cages (Allentown Inc.) for the duration of the experiment. After 8 days, 0.2 mL of *C. difficile* (∼10,000 CFU) was introduced into the mice by oral gavage. Mice were maintained for a total of two weeks after the first colonization with the core community (day 0). Groups of mice (4 mice) with the same core community and *C. difficile* strain were co-housed in a single cage. Mice were weighed and fecal samples were collected at specific time points after oral gavage for NGS sequencing and CFU counting. Cecal contents from mice that were dead or sacrificed were collected for NGS sequencing, CFU counting, and toxin assay.

### Genomic DNA extraction from fecal and cecal samples

The DNA extraction for fecal and cecal samples was performed as described previously with some modifications ^132^. Fecal samples (∼50 mg) were transferred into solvent-resistant screw-cap tubes (Sarstedt Inc) with 500 μL 0.1 mm zirconia/silica beads (BioSpec Products) and one 3.2 mm stainless steel bead (BioSpec Products). The samples were resuspended in 500 μL of Buffer A (200 mM NaCl (DOT Scientific), 20 mM EDTA (Sigma) and 200 mM Tris·HCl pH 8.0 (Research Products International)), 210 μL 20% SDS (Alfa Aesar) and 500 μL phenol/chloroform/isoamyl alcohol (Invitrogen). Cells were lysed by mechanical disruption with a bead-beater (BioSpec Products) for 3 min twice, while being placed on ice for 1 min in between to prevent overheating. Next, cells were centrifuged for 7 min at 8,000 x g at 4°C, and the supernatant was transferred to an Eppendorf tube. We added 60 μL 3M sodium acetate (Sigma) and 600 μL isopropanol (LabChem) to the supernatant and incubated on ice for 1 h. Next, samples were centrifuged for 20 min at 18,000 x g at 4°C, and the supernatant was decanted. The harvested DNA pellets were washed once with 500 μL of 100% ethanol (Koptec), and the remaining trace ethanol was removed by air drying the samples. Finally, the DNA pellets were resuspended into 300 μL of AE buffer (Qiagen). The crude DNA extracts were purified by a Zymo DNA Clean & Concentrator™-5 kit (Zymo Research) prior to PCR amplification and NGS sequencing.

### *C. difficile* colony-forming unit counting from fecal and cecal samples

1. *C. difficile* selective plates were prepared by autoclaving *C. difficile* agar (Oxoid CM0601) and adding defibrinated horse blood (Lampire 7233401, 70 mL/1L media), norfloxacin (Santa Cruz 215586, 120 μg/mL), moxalactam (Santa Cruz 250419, 320 μg/mL), and erythromycin (Santa Cruz 204742, 100 μg/mL) after the media is cooled to ∼55°C. Right after mice fecal or cecal collection, around 1μL of fresh fecal samples were taken using an inoculating loop and mixed with PBS. The samples were then serially diluted (1:10 dilution) using PBS. Four dilutions of each sample were spotted on *C. difficile* selective agar plates, with 2 technical replicates per sample. Plates were incubated at 37°C for 48 h at which point colonies were counted in the dilution spot containing between 5 and 100 colonies. The CFU/mL for each sample was calculated as the average of the 2 technical replicates times the dilution factor. The lower limit of detection for the assay was 20,000 CFU/mL.

## Data availability

Whole-genome sequencing data will be deposited to the NCBI database. RNA-seq data used in this study will be deposited in the NCBI database. Raw DNA sequencing data and processed sequencing data to determine community composition will be made available via Zenodo prior to publication. Exo-metabolomics profiling data will be made available via Zenodo prior to publication or provided as Source Data file.

## Code availability

Codes for fitting the logistic growth model, processing sequencing data and fitting the gLV models will be available through Github prior to publication. The codes are provided to the reviewers during peer review.

## Acknowledgment

We would like to thank Tyler D. Ross, Erin Ostrem, and Yu-Yu Cheng for their helpful advice on this project. This research was supported by the National Institutes of Allergy and Infectious Diseases under grant number R21AI156438 and R21AI159980 for O.S.V, R35GM124774 for O.S.V. Dr. Simcox is an HHMI Freeman Hrabowski Scholar. Research reported in this publication was supported by the Glenn Foundation and American Federation for Aging Research (A22068 for J.S.), an R01 through NIH/NIDDK (R01DK133479 for J.S.), and the Dr. Stephen Babcock AG Chem Fellowship (for I.J.). The funders had no role in study design, data collection and analysis, decision to publish or preparation of the manuscript.

## Authors contribution

J.E.S. and O.S.V. conceived the study. J.E.S. carried out the experiments. J.T., P.L.K.C., and Y.Q. implemented computational modeling for the logistic growth model and gLV models. J.M., I.J., and J.S. performed LC-MS metabolomics of bacterial supernatants. J.E.S. and O.S.V. analyzed data. J.E.S. and O.S.V. wrote the paper and all authors provided feedback on the manuscript. O.S.V. secured funding.

