## Supplementary Figures for "Human gut microbiota interactions shape the long-term growth dynamics and evolutionary adaptations of *Clostridioides difficile*"

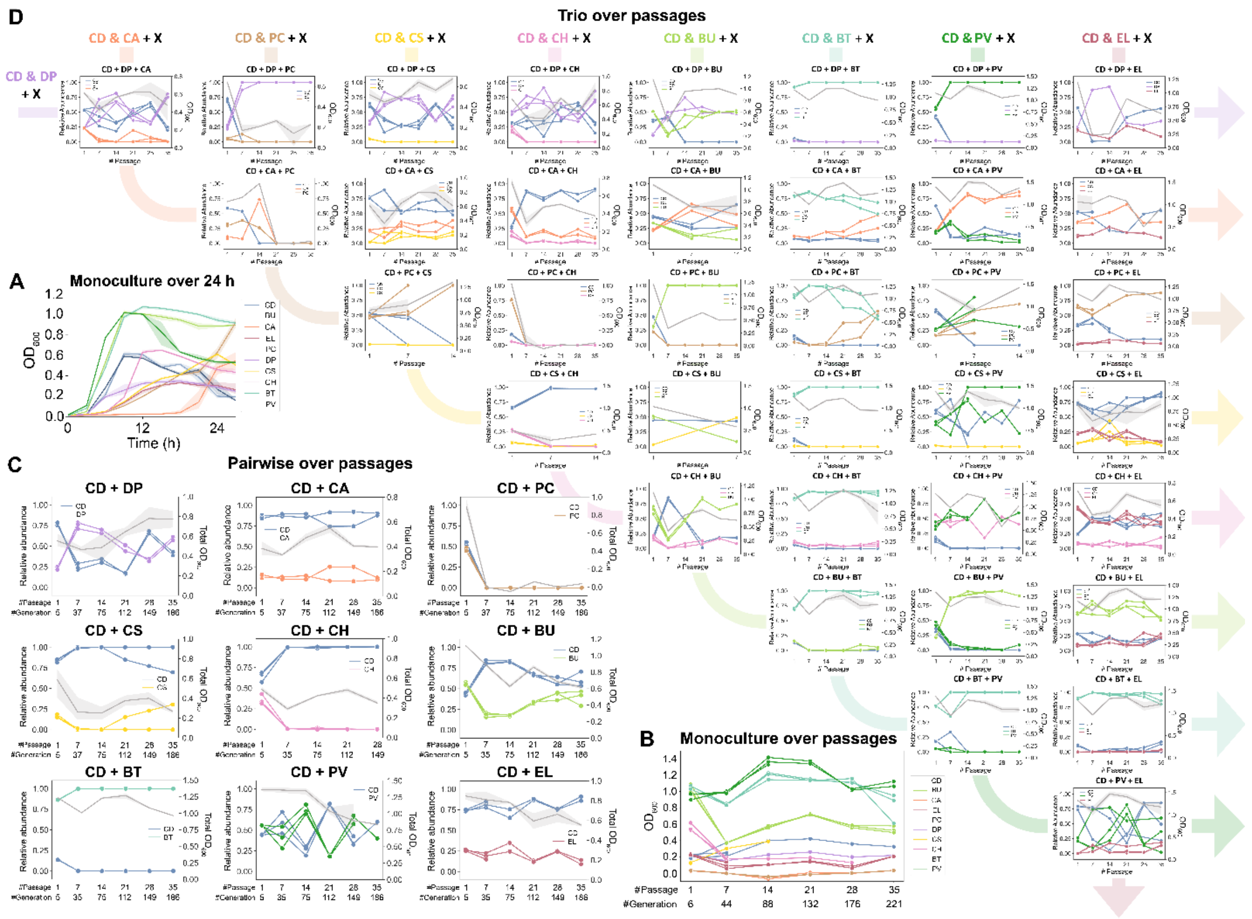


**Supplementary Figure 1. Growth of monocultures, pairwise, and three-member communities consisting of *C. difficile* and different gut species for 35 passages. a**, Absolute abundance ($OD_{600}$) of *C. difficile* and gut bacteria grown in Defined Media (DM) measured over 27 h. Data were shown as mean and 95% c.i. (shading), n = 3. **b**, Absolute abundance ($OD_{600}$) of *C. difficile* and gut bacteria grown for 35 passages in monoculture (n=3). **c-d**, Community dynamics of *C. difficile* and gut bacteria in pairwise (**c**) and three-member communities (**d**) over 35 passages. Dots connected by colored lines indicate the relative abundance of each species (left y-axis) whereas the grey line with shaded 95% confidence interval (CI) indicates the $OD_{600}$ of the community (right y-axis) (n=3). Data from wells that are cross-contaminated were not shown, thus several data points have shorter endpoints than the others.

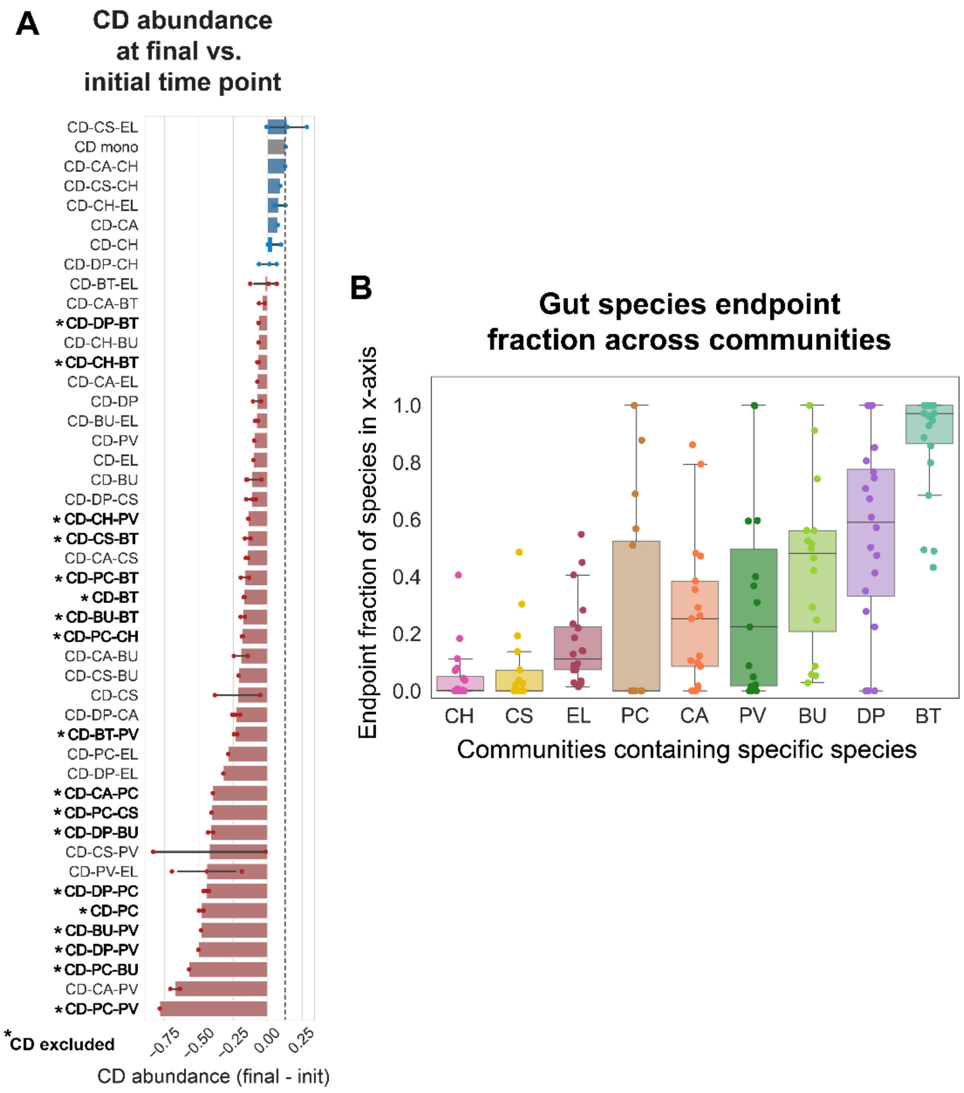


**Supplementary Figure 2. *C. difficile* abundance after long-term passaging with the human gut species. a**, Difference in *C. difficile* absolute abundance at the final timepoint compared to the initial timepoint across all pairwise and three-member communities. Data were shown as mean ± s.d. (n=3) with individual data points. Blue bars indicate an increase in *C. difficile* abundance whereas red bars indicate a decrease in *C. difficile* abundance. Grey bar and the horizontal dashed line mark the change in abundance of *C. difficile* monoculture. Communities that are marked with asterisks are those where *C. difficile* was excluded from the community. **b**, Fraction of individual gut species at the last time point of all pairwise and three-member communities extracted from **Fig. S1c-d**.

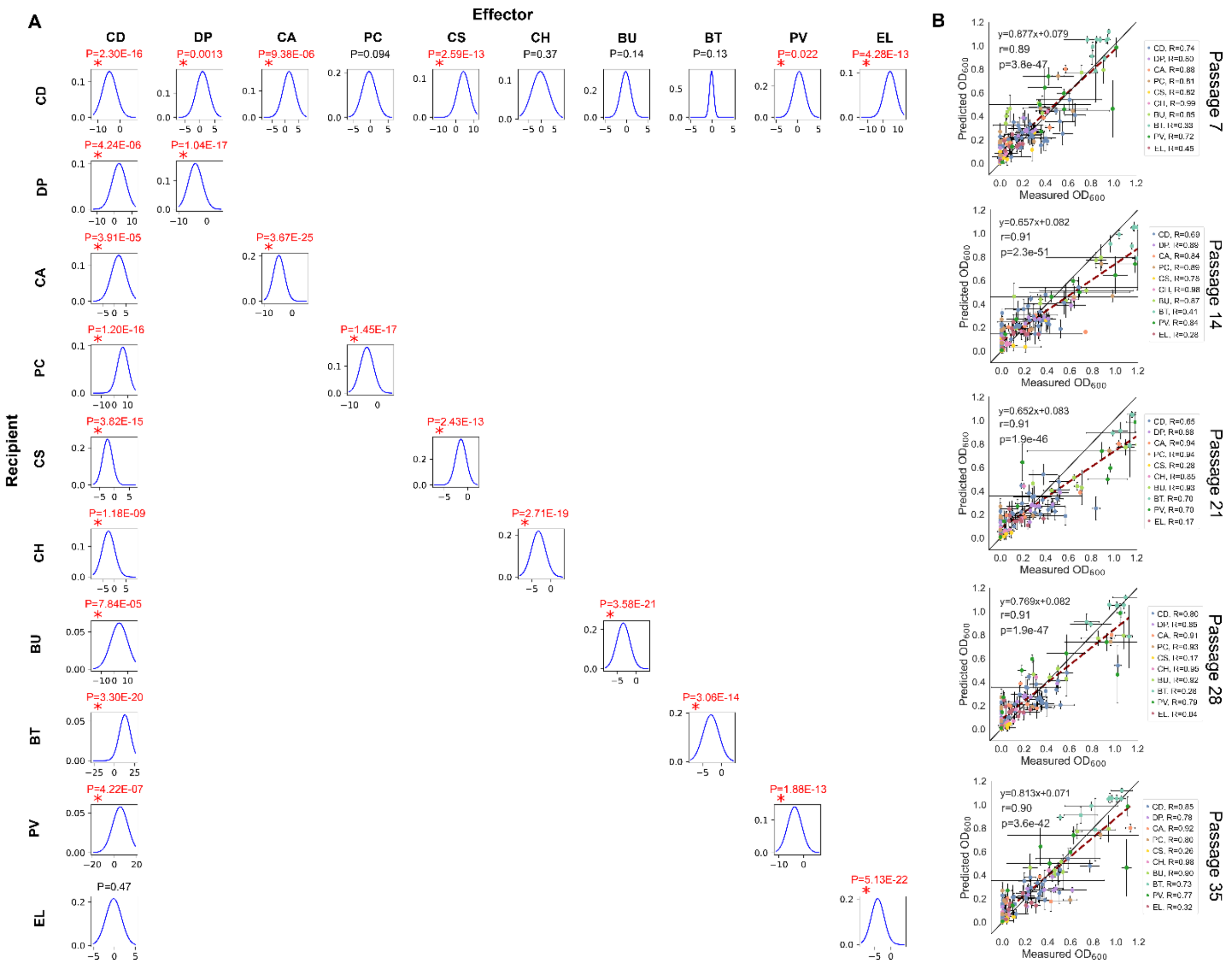


**Supplementary Figure 3. Parameter estimates for the gLV model fit to the long-term passaging data over 35 passages. a**, The plots show the mean and variance of each gLV parameter ($a_{ij}$ values, see **Methods**). Data used to fit the gLV model is from DATASET001 (**Table S3**). Asterisks indicate parameters where the means significantly deviate from 0 based on unpaired *t*-test (two-sided). **b**, Scatter plot of gLV model fit versus measured species absolute abundances at each passage. Horizontal error bars indicate standard deviation of measured $OD_{600}$ (n=1-3). Vertical error bars indicate the standard deviation calculated from 12 models out of the 64 models that succeeded (see **Methods**). Colors indicate the species in the community. Red dashed line indicates the linear regression between the mean measured $OD_{600}$ and the predicted $OD_{600}$. Two-sided Pearson's correlation coefficient (*R*) and *p*-values are shown, which was computed using the pearsonr from the scipy package in Python.

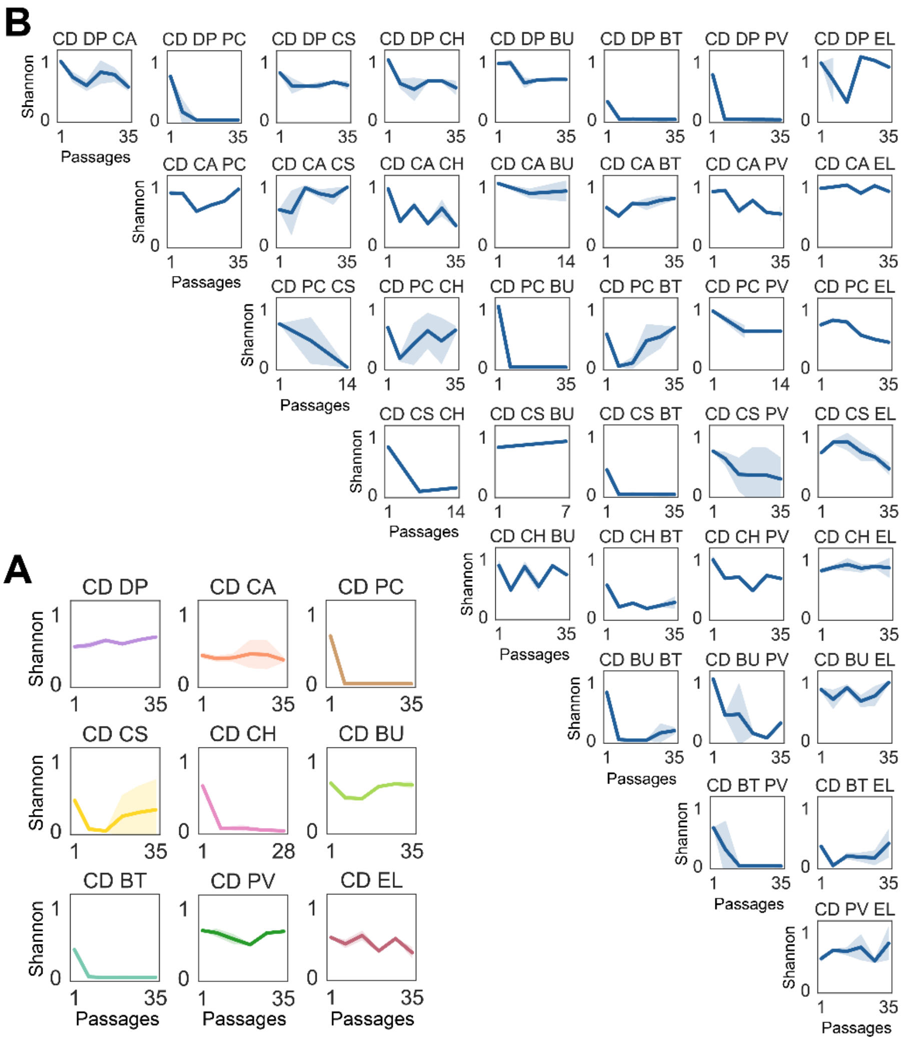


**Supplementary Figure 4. Shannon diversity of the communities throughout the passages. a-b**, Shannon diversity of pairwise (**a**) and three-member (**b**) communities over 35 passages, extracted from **Fig. S1c-d**. Data were shown as mean and 95% c.i. (shading), n = 3.

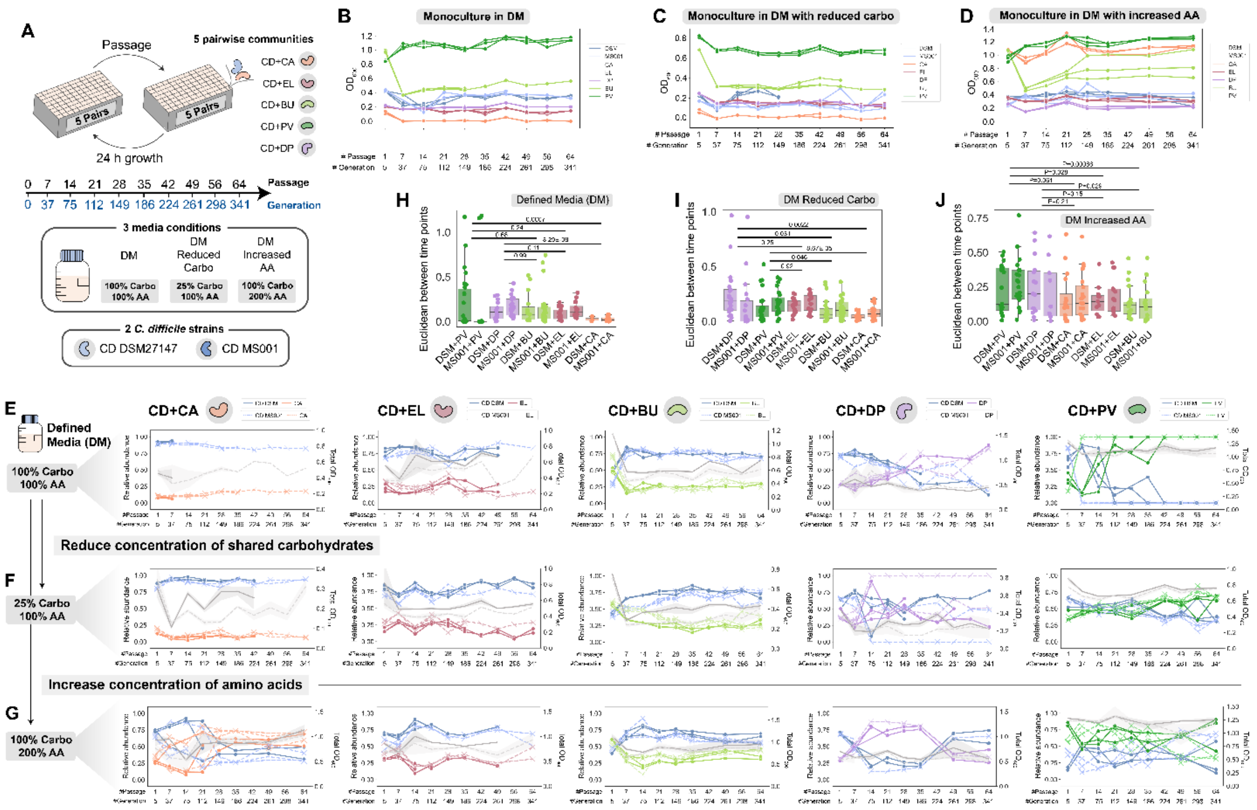


**Supplementary Figure 5. Long-term growth experiment in 3 different media with 2 different *C. difficile* strains over 64 passages. a**, Schematic of the long-term growth experiment of 5 pairwise communities over 64 passages. The communities were grown using two *C. difficile* strains (DSM 27147 and MS001) and three media conditions (Defined Media (DM), DM with 75% less carbohydrate concentrations, and DM with 100% higher amino acid concentrations). **b-d,** Monoculture absolute abundance ($OD_{600}$) of *C. difficile* DSM27147 and MS001 and 5 gut bacteria grown over 64 passages (~341 generations) in DM (**b**), DM with reduced carbohydrates (**c**), and in DM increased amino acids (**d**) (n=3). DSM27147 is the hypervirulent R20291 reference strain of the epidemic ribotype 027. MS001 is a clinical isolate that possesses a larger genome with a higher number of mobile elements including conjugative systems, plasmids, and phages compared to DSM27147. **e-g**, Growth of pairwise communities consisting of *C. difficile* DSM 27147 strain (solid lines) or *C. difficile* MS001 strain (dashed lines) with one of 5 gut species over 341 generations in DM (**e**), DM with reduced carbohydrates (**f**), and in DM with increased amino acids (**g**) (n=3). Dots connected by colored lines indicate the relative abundance of each species (left y-axis) as measured by 16S rRNA sequencing, whereas the grey line with shaded 95% confidence interval (CI) indicates the $OD_{600}$ of the community (right y-axis). Data from contaminated wells are excluded, thus several data points have shorter endpoints than the others. **h-j**, Box plots of Euclidean distances between time point measurements throughout 64 passages in all pairwise communities in DM (**h**), in DM with reduced carbohydrates (**i**), and in DM with increased amino acids (**j**). *p*-values from unpaired *t*-test (two-sided) of the Euclidean distances between two pairwise communities containing *C. difficile* are shown (data from *C. difficile* DSM and MS001 are combined).

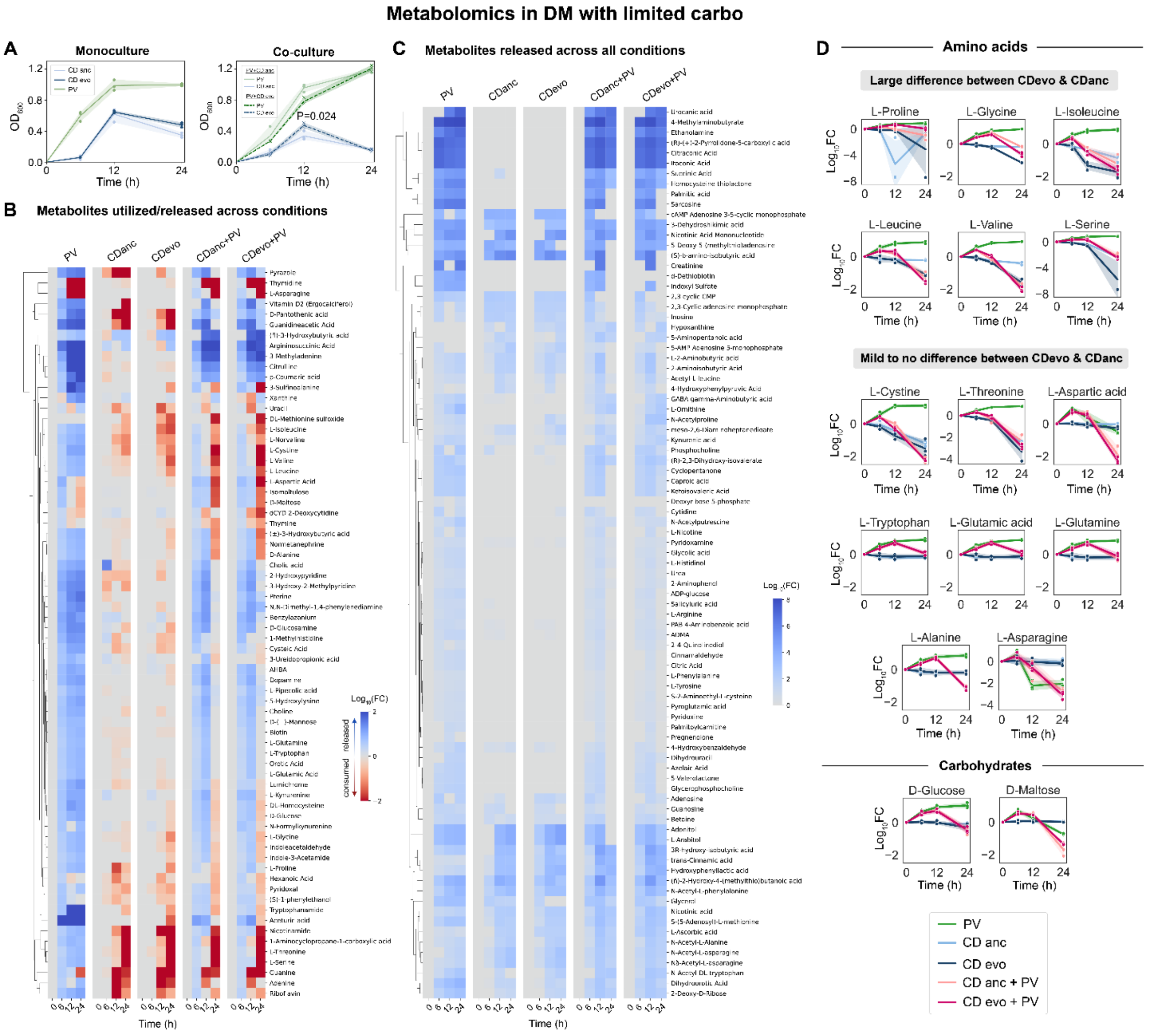


**Supplementary Figure 6. Exo-metabolomic profiling of *C. difficile* and *P. vulgatus* in DM with limited carbohydrates. a,** Absolute abundance of the species over time. For monocultures, species absolute abundance was quantified using $OD_{600}$. For co-cultures, species absolute abundance was quantified by multiplying the relative abundance from 16S sequencing with the total community $OD_{600}$. Data were shown as mean and s.d. (shading), n = 3. Significant *p*-value from unpaired *t*-test (two-sided) of the absolute abundance between the evolved and the ancestral *C. difficile* strain is shown. **b-c**, Heatmap of fold change of the metabolites utilized and released by PV, ancestral *C. difficile,* and evolved *C. difficile* strains in monoculture and co-culture compared to the blank media (**b**) and heatmap of fold change of the metabolites released across PV, ancestral *C. difficile,* and evolved *C. difficile* strains monoculture and co-culture compared to the blank media (**c**). Cultures were grown in DM with reduced carbohydrate concentration for 24 h. At 0, 6, 12, and 24 h, supernatants were subjected to LC/MS. All non-zero values in the heatmap are statistically significant (Two-sided *t*-test with unequal variance). **d**, Utilization/release dynamics of amino acids and carbohydrates in PV, ancestral *C. difficile,* and evolved *C. difficile* strains in monocultures and co-cultures. Data were shown as mean and s.d. (shading), n = 3.

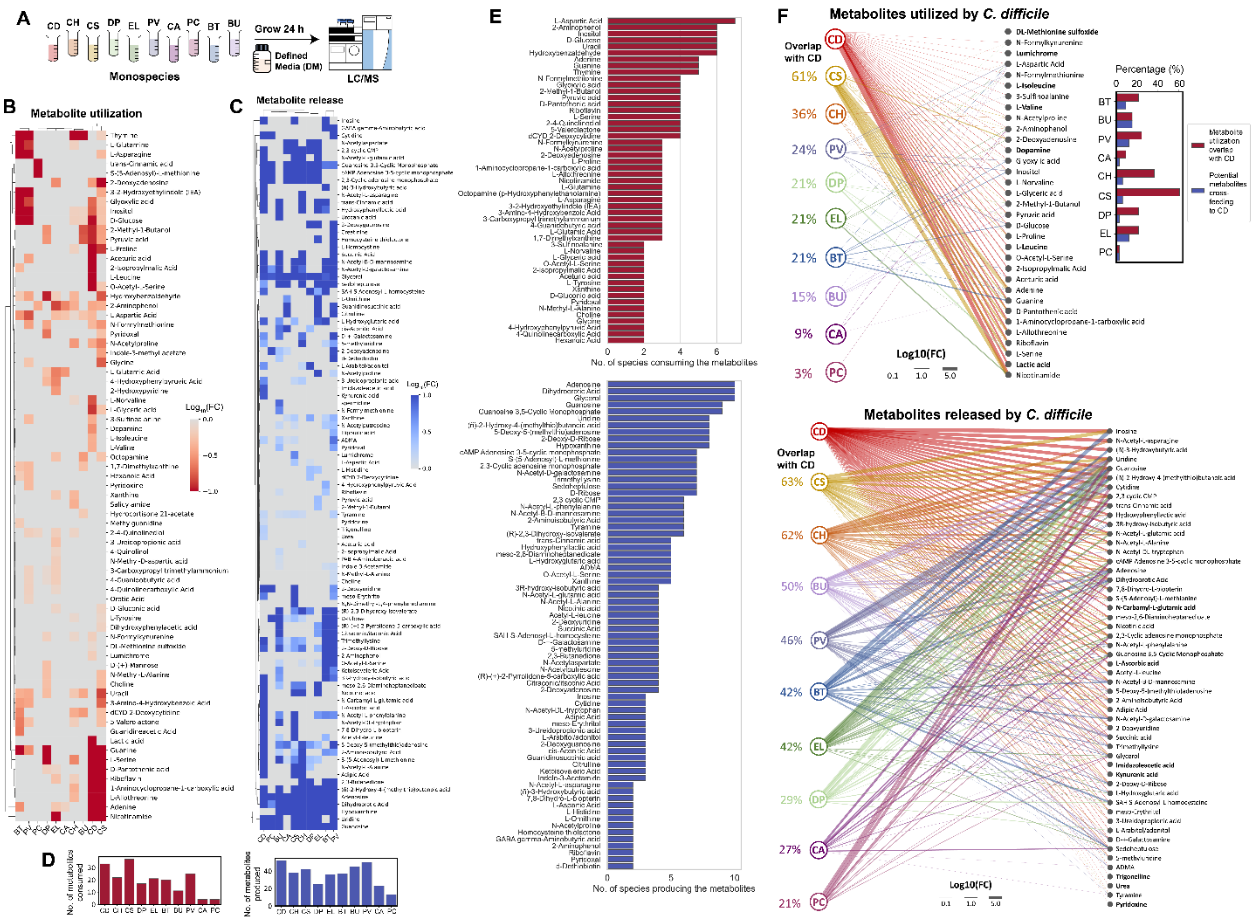

**Supplementary Figure 7. Exo-metabolomic profiling of *C. difficile* and 9 gut bacteria in DM.** **a,** Exo-metabolomic profiling of *C. difficile* and 9 gut bacteria. Individual bacteria were grown in DM for 24 h and the supernatant was subjected to LC/MS analysis (n=3). **b-c,** Heatmap of fold change of the metabolites utilized (**b**) and released (**c**) by *C. difficile* and the 9 gut species compared to the blank media. Red/blue colors were those that have statistically higher or lower concentrations compared to the blank media control (Two-sided *t*-test with unequal variance). **d**, Number of metabolites consumed (**left**) or released (**right**) by each of the species. **e**, Number of species consuming (**top**) or releasing (**bottom**) specific metabolites. Only metabolites that were consumed or released by more than one species were shown. **f**, Bipartite networks of metabolite utilization (**top**) or release (**bottom**) profile of *C. difficile* and the 9 gut bacteria. Only metabolites utilized/released by *C. difficile* are shown. The complete metabolite utilization/release profile is shown in **panel b-c**. Only metabolites that have statistically lower/higher concentrations compared to the blank media control are shown (Two-sided t-test with unequal variance). Metabolites in bold were uniquely utilized/released by *C. difficile*. Edges represent $\log_{10}$ fold change of metabolite utilization/release. The degree of overlap in metabolite utilization/release compared to *C. difficile* is calculated from the number of edges coming out of a particular species divided by the number of edges coming out of *C. difficile*. The inset figure on the top right shows the % of potential metabolite cross feeding versus competition with *C. difficile*. Potential cross-feeding to *C. difficile* is defined as those being consumed by *C. difficile* but produced by the other gut species. Competition or utilization overlap is defined as those being consumed by both *C. difficile* and the other gut species.

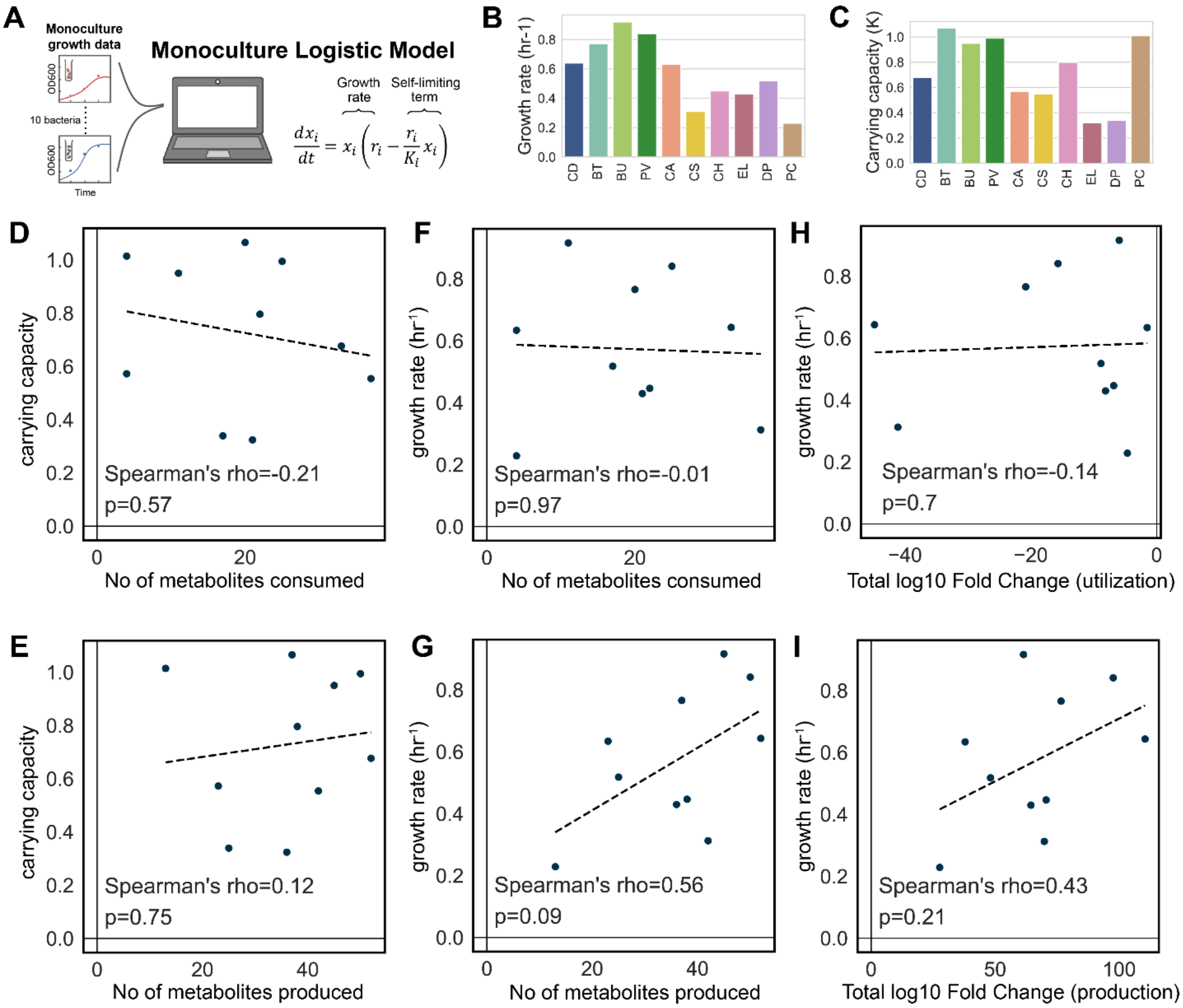


**Supplementary Figure 8. Monoculture growth rate and carrying capacity of *C. difficile* and the 9 gut bacteria in DM. a,** Monoculture growth data of *C. difficile* and 9 gut bacteria in DM (**Fig. S1a**) were fitted to the logistic model (See **Methods**). Mathematical description of the logistic growth model was shown, where $x_i$ is the absolute abundance of species $i$, parameter $r_i$ is its maximum growth rate, and $K_i$ is its steady-state abundance or carrying capacity. When fitting the experimental data to the model, we cut timepoints where $OD_{600}$ drops above 10% to exclude the death phase. **b-c**, Growth rate (**b**) and carrying capacity (**c**) for the 10 bacteria from the model fitting. **d-e**, Correlations between carrying capacity and the number of metabolites consumed (**d**) and released (**e**). **f-g**, Correlations between growth rate and the number of metabolites consumed (**f**) and released (**g**). **h-i**, Correlations between growth rate and the total log10 fold change in metabolite utilization (**h**) and release (**i**). Two-sided Spearman's rho and *p*-value are shown, which was computed using the spearmanr from the scipy package in Python.

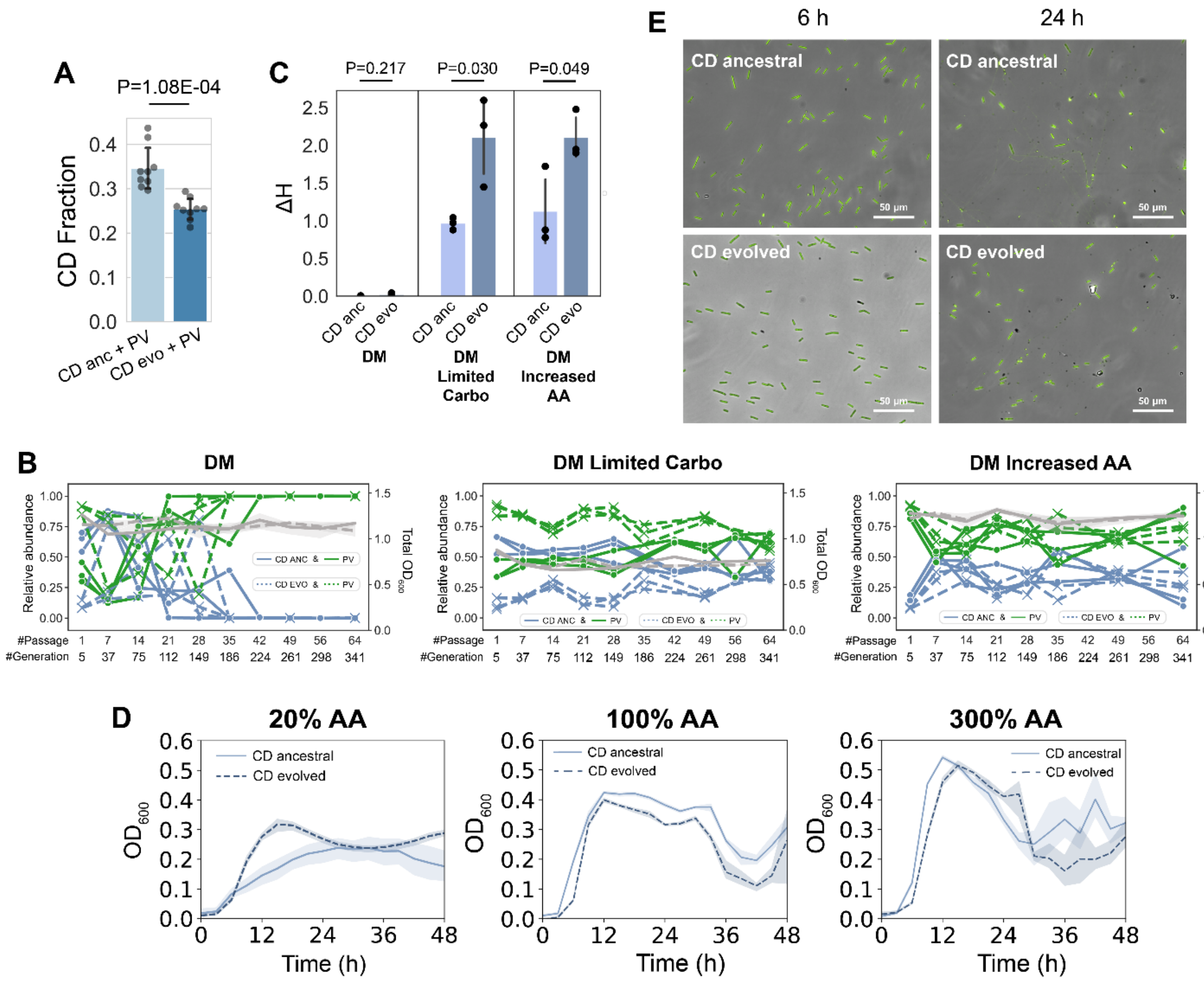


**Supplementary Figure 9. Growth characterization of the evolved *C. difficile* strain compared to the ancestral strain**. **a**, Bar plots of the composition of evolved and ancestral *C. difficile* strains when grown with PV in DM with reduced carbohydrates (mean ± s.d., n=9). The *p*-value from unpaired *t*-test (two-sided) is shown. **b**, Growth dynamics of the evolved *C. difficile* strain with PV over 64 passages (~341 generations) in DM (**left**), DM with reduced carbohydrates (**middle**), and in DM with increased amino acids (**right**) (dashed lines, n=3). The growth dynamics of the ancestral *C. difficile* strain with PV from **Fig. S5e-g** were shown for comparison (solid lines). Markers connected by colored lines indicate the relative abundance of each species (left y-axis) as measured by 16S rRNA sequencing, whereas the grey line with shaded 95% confidence interval (c.i.) indicates the $OD_{600}$ of the community (right y-axis). **c**, Degree of coexistence (ΔH, measured by the Shannon diversity at the last time point relative to the initial time point) in CD-PV co-cultures containing the ancestral or evolved *C. difficile* strain across different media conditions (mean ± s.d., n=3). The *p*-values from unpaired *t*-test (two-sided) are shown. **d**, Monoculture growth ($OD_{600}$) of ancestral and evolved *C. difficile* strains in DM supplemented with 20% (left), 100% (middle), and 300% (right) amino acid concentrations. Data were shown as mean and 95% c.i. (shading), n = 3. Solid lines indicate ancestral *C. difficile* strain whereas dashed lines indicate evolved *C. difficile* strain. **e**, Fluorescence microscopy of the ancestral and evolved *C. difficile* strains grown anaerobically in DM. The cultures were sampled at two time points (6 h and 24 h) and stained with SYBR Green dye.

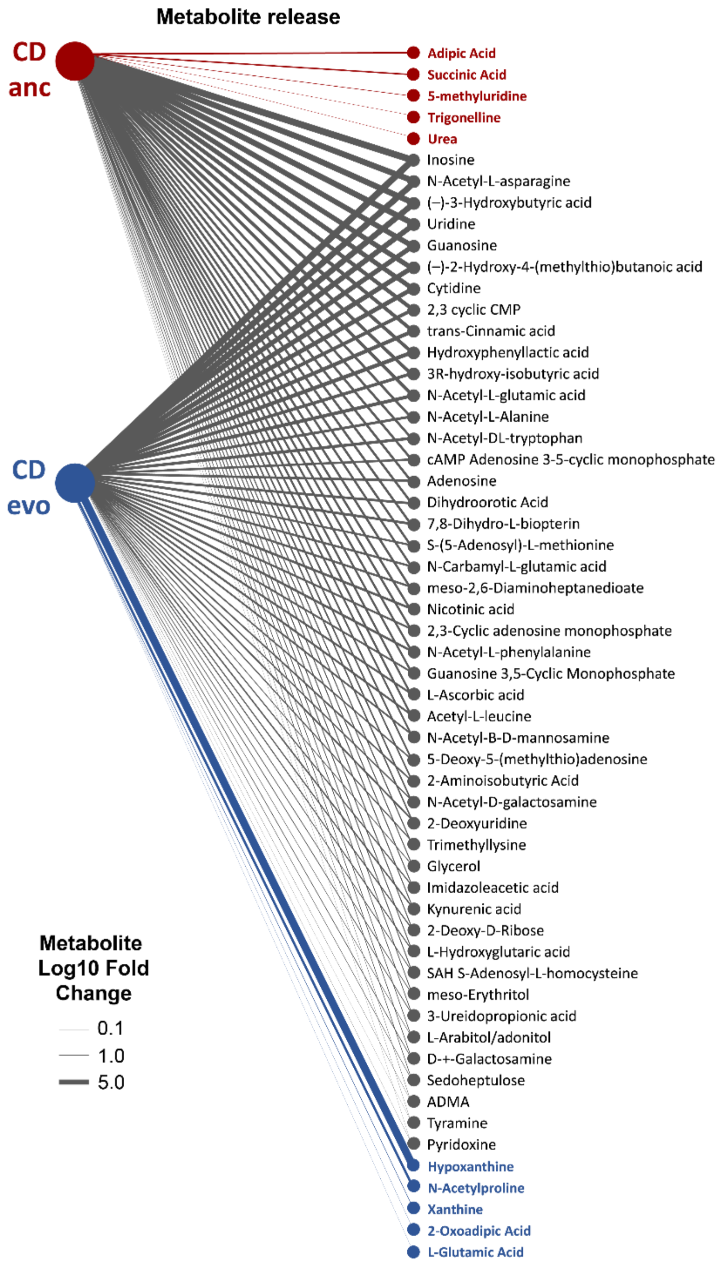


**Supplementary Figure 10. Metabolite release profile of the evolved *C. difficile* strain compared to the ancestral strain in DM.** Bipartite network of metabolite release between the ancestral and evolved *C. difficile* strains. *C. difficile* strains were grown in DM for 24 h and the supernatant was subjected to LC/MS analysis. Three biological replicates were performed for each strain, and metabolites that have significantly higher concentrations compared to the blank media are shown (two-sided *t*-test with unequal variance). Metabolites bolded with red (blue) are uniquely released by the ancestral strain (evolved strain). Metabolites marked with red (blue) asterisks have >10-fold higher release in the ancestral strain (evolved strain). Edges represent $\log_{10}$ fold change of metabolite release.

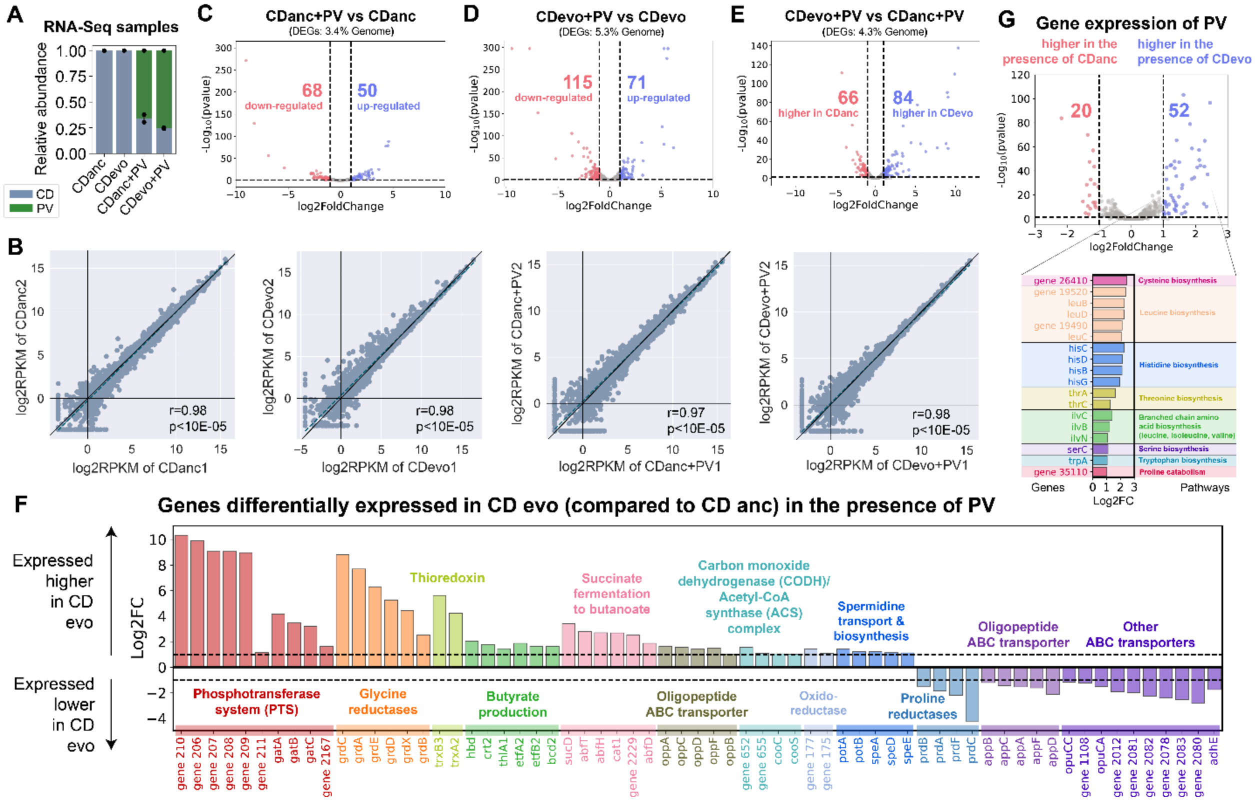


**Supplementary Figure 11. Transcriptomic profiling of the evolved *C. difficile* strain and the ancestral strain. a,** Stacked bar plots show the composition of the samples subjected to RNA-Seq as determined by 16S sequencing. Each bar represents the average relative abundance of each species, and the error bars represent s.d. (n=2). **b**, Scatter plot showing the correlation between the Reads Per Kilobase Million (RPKM) of the two biological replicates in each sample. Blue dashed line indicates the linear regression between the coefficients of the two communities. Two-sided Pearson's correlation coefficient ($r$) and $p$-values are shown. **c-e**, Volcano plots of log-transformed transcriptional fold changes for ancestral *C. difficile* strain in the presence of PV compared to monoculture (**c**), evolved *C. difficile* strain in the presence of PV compared to monoculture (**d**), and evolved *C. difficile* strain compared to ancestral *C. difficile* strain in the presence of PV (**e**). Vertical dashed lines indicate 2-fold change, and the horizontal dashed line indicates the statistical significance threshold as calculated by DESeq2's Wald test with Benjamini-Hochberg multiple testing correction (BH-adjusted p value=0.05). Blue indicates up-regulated genes and red indicates down-regulated genes. **f**, Bar plot of the log-transformed fold changes of selected genes that were expressed higher and lower in the evolved *C. difficile* strain compared to the ancestral *C. difficile* strain in the presence of PV. Horizontal dashed lines indicate a 2-fold change. **g**, DEGs in PV in the presence of evolved *C. difficile* strain compared to ancestral strain. The volcano plot shows the log-transformed transcriptional fold changes for PV in the presence of evolved *C. difficile* strain compared to ancestral strain. The bar plot shows the log-transformed fold changes of selected genes that were expressed higher in PV in the presence of evolved *C. difficile* strain compared to ancestral strain.

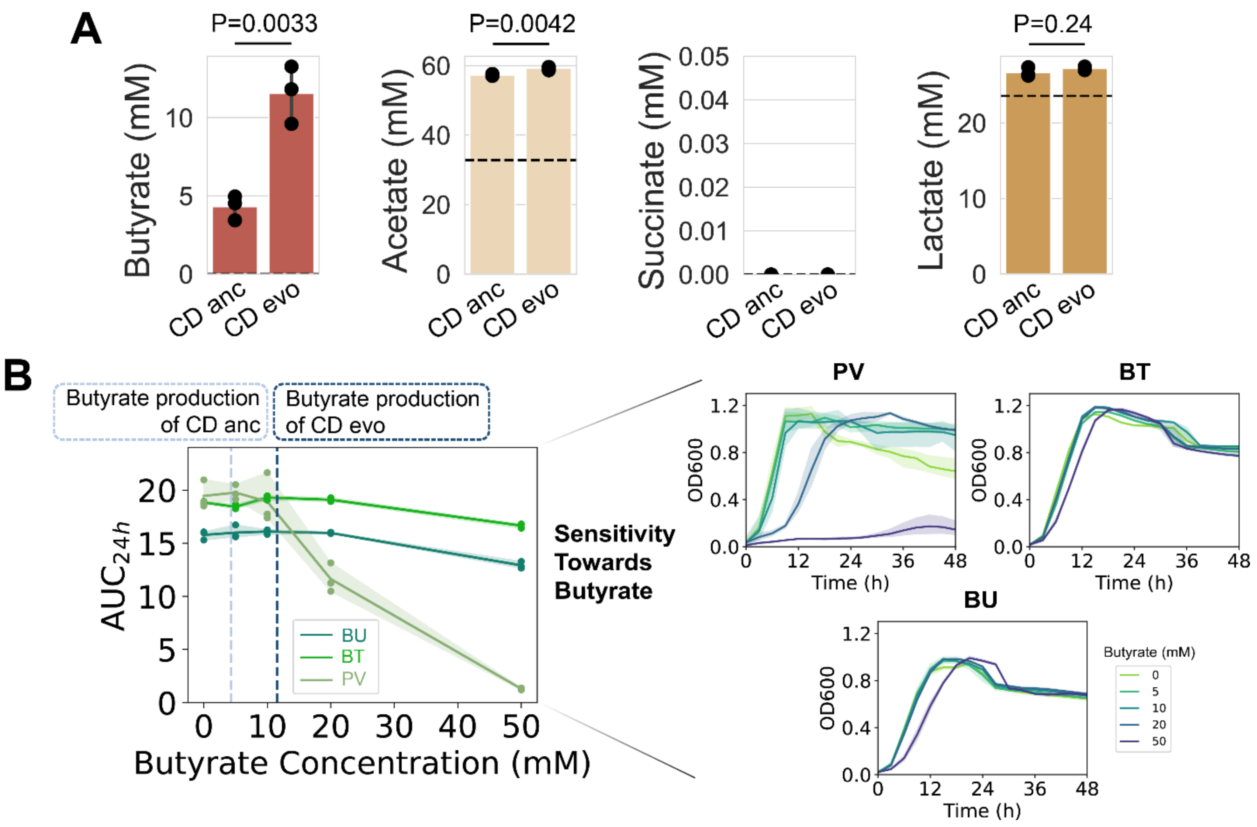


**Supplementary Figure 12. Production of butyrate, acetate, succinate, and lactate by the evolved and ancestral *C. difficile* strains. a**, Quantification of organic acids in ancestral *C. difficile* and evolved *C. difficile* strains in DM (mean ± s.d., n=3). Horizontal dashed lines indicate the concentration detected in the blank media. *p*-values from unpaired *t*-test (two-sided) are shown. **b**, Effects of butyrate on the growth of *Bacteroides* species. Left panel shows the sensitivity of *Bacteroides* species monoculture growth towards butyrate concentration in DM. Mean (line) and 95% c.i. (shading) with individual data points were shown (n = 3). Vertical dashed lines indicate the butyrate production level of ancestral *C. difficile* (light blue) and evolved *C. difficile* strain (dark blue) in the same media. $AUC_{24h}$ were calculated from the growth curves in the right panel, showing the time-course $OD_{600}$ measurements of PV, BT, and BU in DM with varying concentrations of butyrate. Data were shown as mean and 95% c.i. (shading), n = 3.

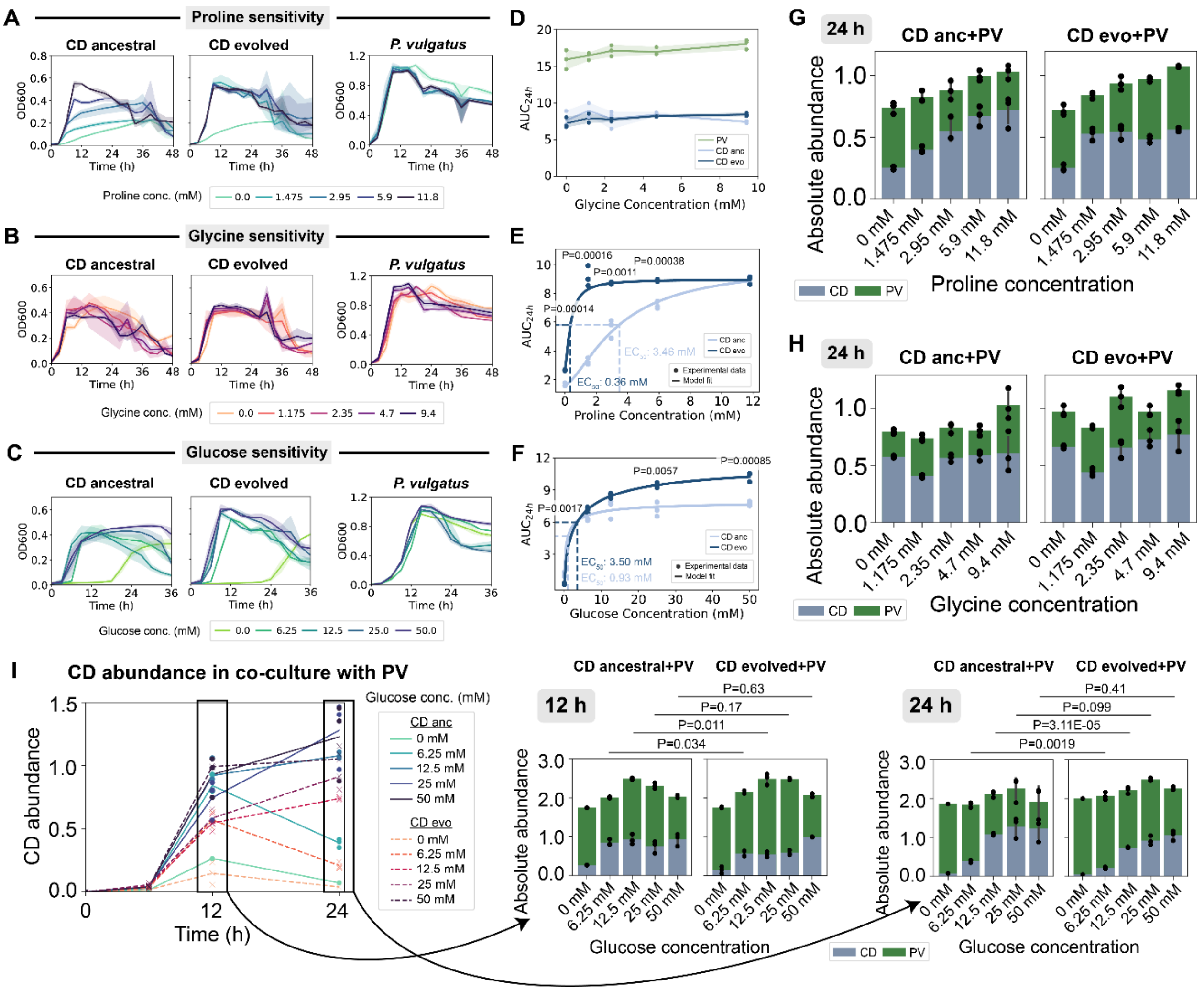


**Supplementary Figure 13. Growth differences between the evolved and ancestral *C. difficile* strain. a-c**, Time-course $OD_{600}$ measurements of the *C. difficile* strains and PV in DM with varying concentrations of proline (**a**), glycine (**b**), and glucose (**c**). Data were shown as mean and 95% c.i. (shading), n = 3. **d**, Growth of evolved and ancestral *C. difficile* strains and PV under different glycine concentrations. $AUC_{24h}$ was calculated from the growth curves in **panel b**. Mean (line) and 95% c.i. (shading) with individual data points are shown (n = 3). **e-f**, Growth of evolved and ancestral *C. difficile* strains under different proline (**e**) or glucose (**f**) concentrations. $AUC_{24h}$ was calculated from the growth curves in **panel a** and **c** (n=3). Lines indicate experimental data fit to the Hill function (See **Methods**). Significant *p*-values from unpaired *t*-test (two-sided) between the AUC of the evolved and ancestral strain at specific proline or glucose concentration are shown. **g-h**, Stacked bar plots of the *C. difficile* strains grown with PV in media supplemented with different proline (**g**) or glycine (**h**) concentrations for 24 h. Each bar represents the average absolute abundance of each species, and the error bars represent s.d. (n=3). Individual data points were shown. **i**, *C. difficile* abundance when grown with PV in media supplemented with different glucose concentrations. Solid lines indicate ancestral *C. difficile* strain when co-cultured with PV whereas dashed lines indicate evolved *C. difficile* strain when co-cultured with PV. Bar plots on the right show *C. difficile* absolute abundance at 12 h and 24 h in the media with varying glucose concentrations (mean ± s.d., n=3). *p*-values from unpaired *t*-test (two-sided) of the absolute abundance between evolved and ancestral *C. difficile* strain at specific glucose concentration are shown.

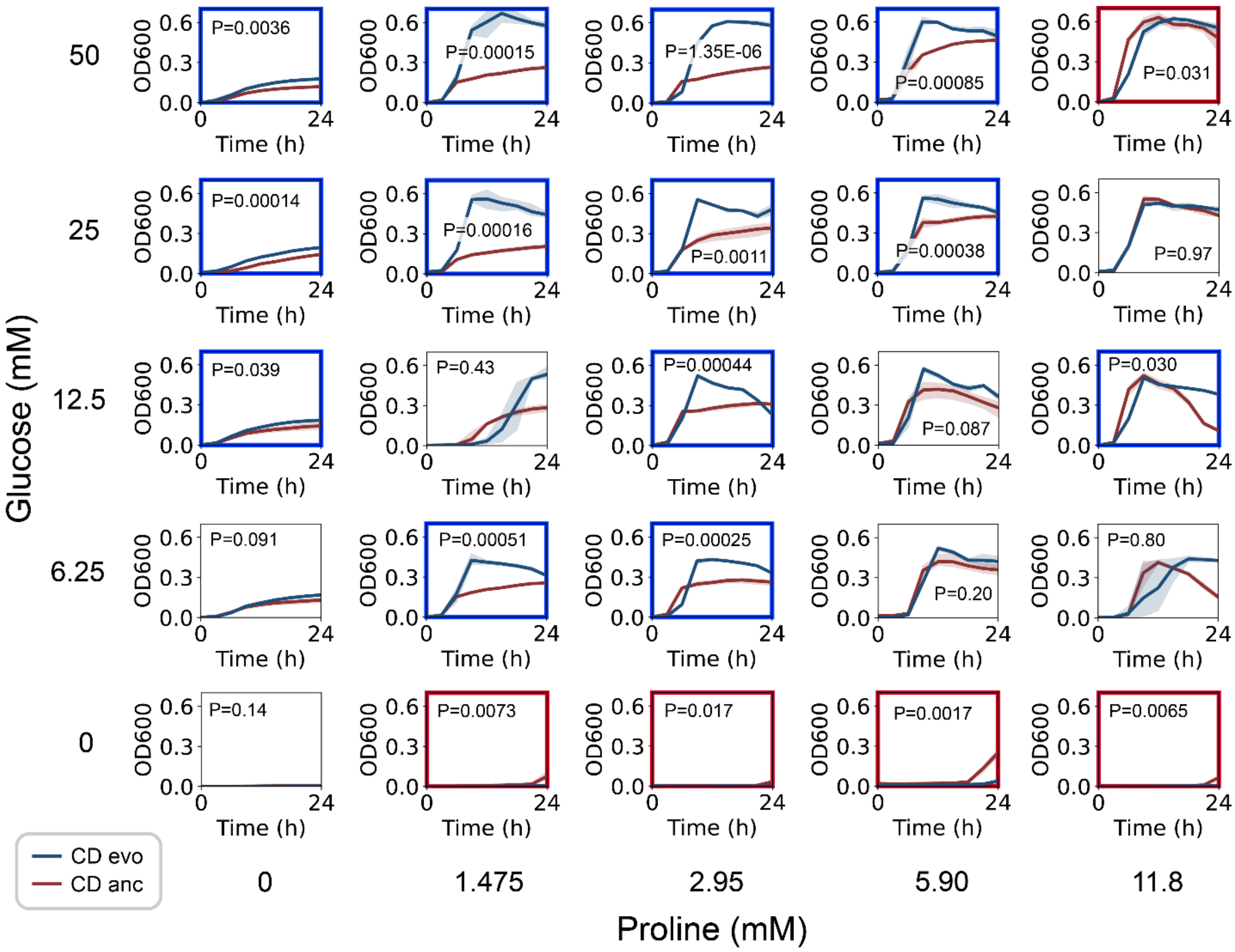


**Supplementary Figure 14. Fitness of the evolved *C. difficile* strain compared to the ancestral strain under different proline and glucose concentrations**. Time-course $OD_{600}$ measurements of the evolved and ancestral *C. difficile* strains under different concentrations of proline and glucose. Data were shown as mean and 95% c.i. (shading), n = 3 biological replicates. Plots outlined with blue (red) indicate those where the $AUC_{24h}$ of the evolved *C. difficile* strain is significantly higher (lower) than the ancestral strain based on unpaired *t*-test (two-sided). The *P*-values are shown within the plot.

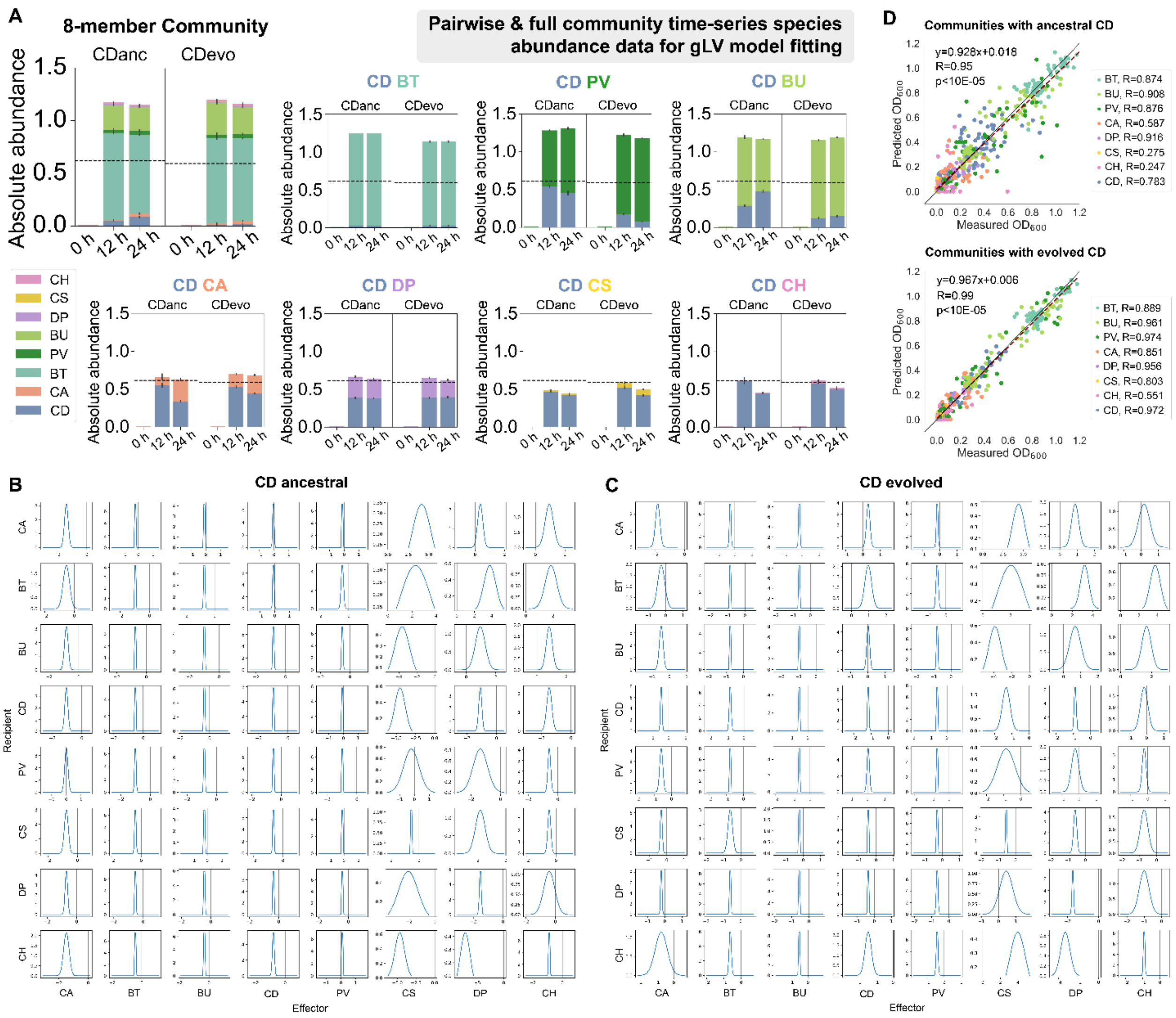


**Supplementary Figure 15. gLV fitting of community growth data containing the ancestral and evolved *C. difficile* strain. a,** Absolute abundance of the full 8-member community and pairwise communities containing either the ancestral or evolved *C. difficile* strain over time when grown in DM. Absolute abundance is the product of 16S relative abundance and community $OD_{600}$. Each bar represents the average absolute abundance of each species, and the error bars represent s.d. (n=3). Horizontal dashed line indicates the $OD_{600}$ of the ancestral and evolved *C. difficile* strain monoculture. **b-c**, Parameter estimates for the gLV model containing the ancestral *C. difficile* strain (**b**) and evolved *C. difficile* strain (**c**). The plots show the mean and variance of each gLV parameter ($a_{ij}$ values) for communities containing the *C. difficile* ancestral and evolved strains (details can be found in the **Supplementary text**). Data used to fit the gLV model is from DATASET002 (**Table S3**). Based on the posterior parameter distributions, we analyzed parameters with absolute values that were significantly constrained to be non-zero based on the Wald test [1]. The Wald test compares the parameter mean to its standard deviation to evaluate whether the peak of the parameter distribution is significantly higher or lower than zero compared to the width of the distribution. The percentage of constrained parameters is 91% and 86% for communities containing the ancestral and evolved *C. difficile* strain respectively. **d,** Model performance for communities containing ancestral and evolved *C. difficile* strains. Scatter plot of measured $OD_{600}$ versus predicted $OD_{600}$ for 2-8 species communities. Colors indicate the species in the community that was measured and predicted. Red dashed line indicates the linear regression between the mean measured $OD_{600}$ and the predicted $OD_{600}$. Two-sided Pearson's correlation coefficient ($R$) and $p$-values are shown, which was computed using the pearsonr from the scipy package in Python.

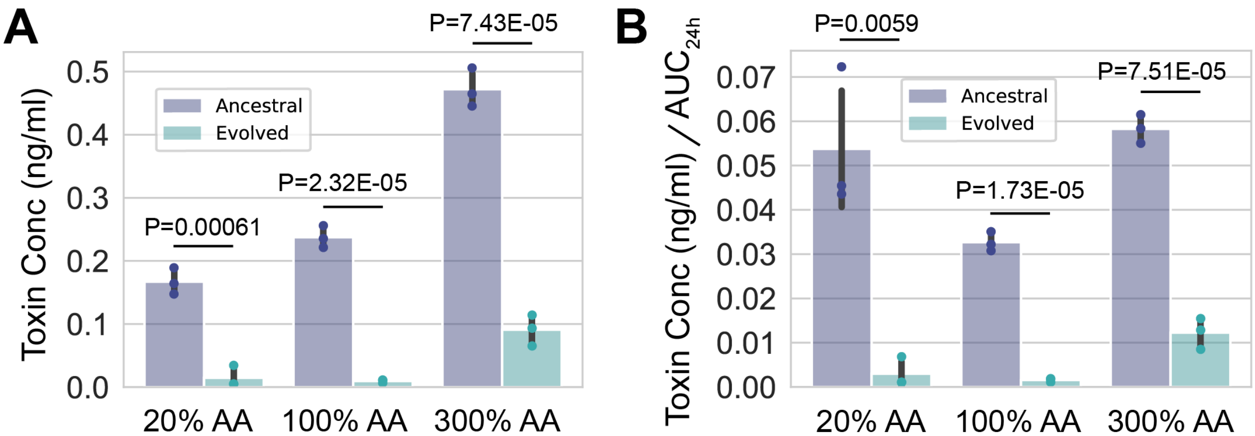


**Supplementary Figure 16. Toxin production and toxin yield of the evolved *C. difficile* strain compared to the ancestral strain. a-b,** Toxin production (**a**) or toxin production per growth of *C. difficile* monocultures (toxin yield) (**b**) in DM containing different amino acid concentrations. Toxin concentrations (TcdA and TcdB) were measured after 24 h of growth using ELISA (mean ± s.d., n=3). In **panel b**, toxin concentration values were normalized with the area under the curve after 24h of growth ($AUC_{24h}$) in the same media from **Fig. S9d**. Bars on the left (blue) show the toxin production/yield of the ancestral *C. difficile* strain whereas bars on the right (green) show the toxin production/yield of the evolved *C. difficile* strain. *p*-values from unpaired *t*-test (two-sided) of the toxin production/yield between evolved and ancestral *C. difficile* strains are shown.

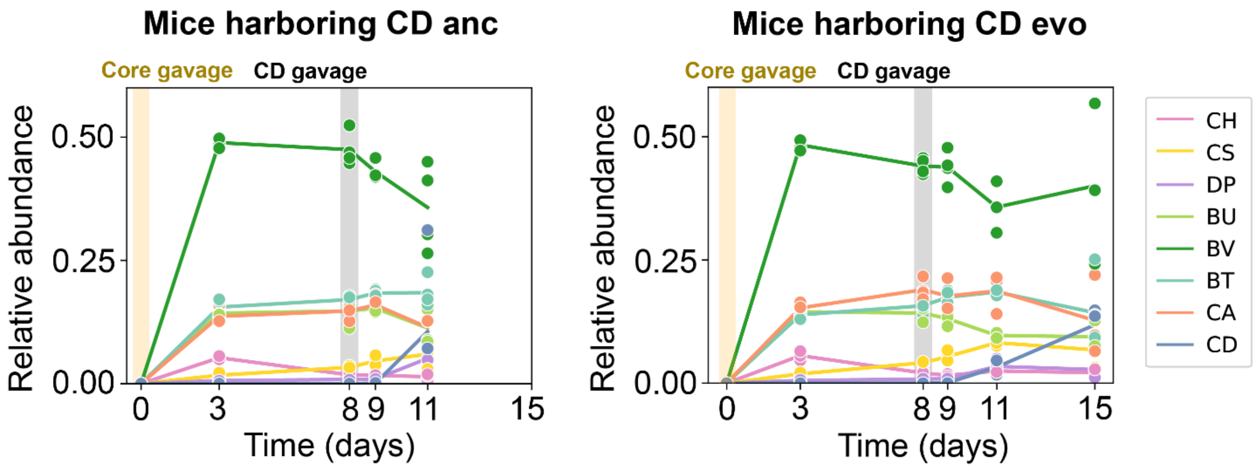


**Supplementary Figure 17. Composition of species in mice gavaged with ancestral and evolved *C. difficile* over time.** Relative abundance was determined using 16S sequencing of mice fecal content (for mice that survived) and cecal content (for dead mice). Data points represent individual mice, and the line represents the average of all mice in the group.

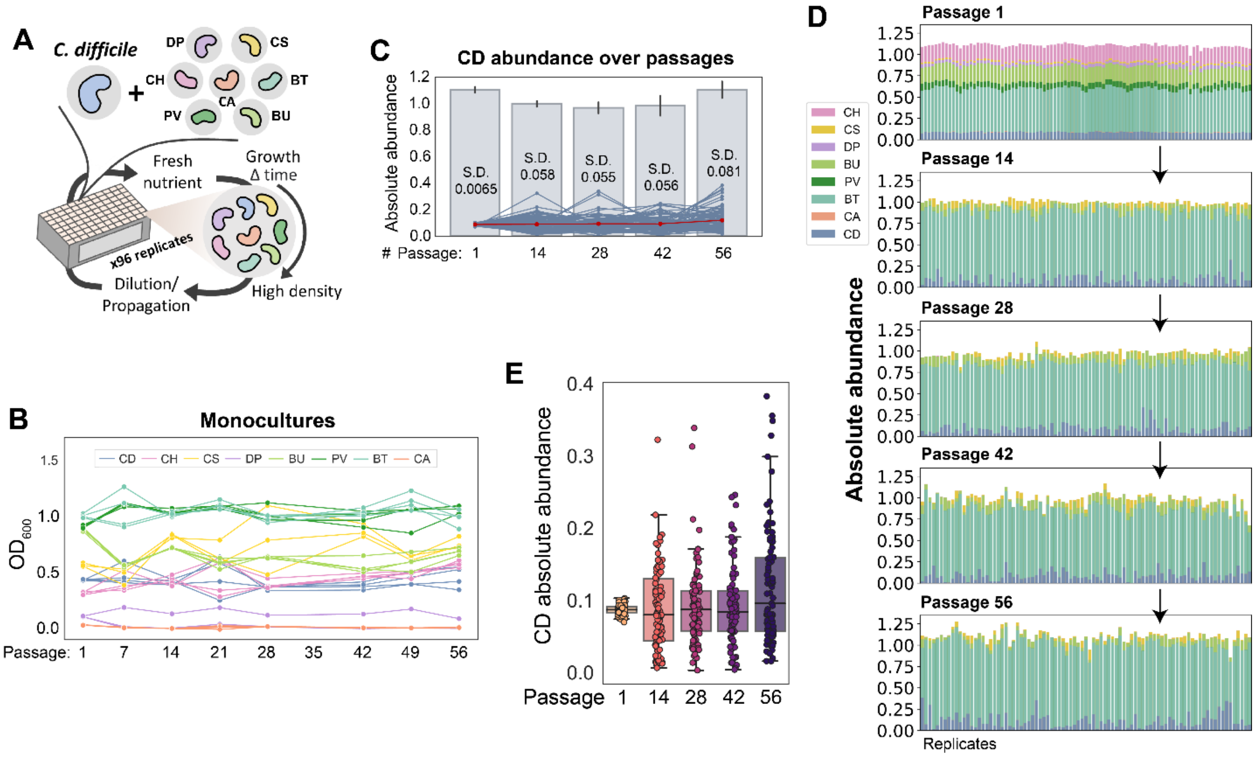


**Supplementary Figure 18. Assessing variability in long-term community assembly containing *C. difficile*. a,** Schematic of simple laboratory passaging experiments to study variability in long-term community assembly. *C. difficile* is grown with the 7 gut species used to study interspecies interactions with 96 independent cultures. To ensure that *C. difficile* stays in the community, the media is supplemented with carbohydrates that are preferred by *C. difficile*: Glucose, sorbitol, mannitol, trehalose, and succinate at a concentration of 2 g/L each. **b**, Monoculture absolute abundance ($OD_{600}$) of *C. difficile* and 7 gut bacteria grown over 56 passages (n=3). **c**, Growth dynamics of *C. difficile* over 56 passages (96 biological replicates). Blue lines indicate the absolute abundance of *C. difficile* from individual replicates, whereas the red line indicates the mean of *C. difficile* abundance over time. Bar plots show the total $OD_{600}$ of the 8-member community over time (mean ± s.d., n=96). The s.d. values of *C. difficile* absolute abundance across the 96 replicates for each passage is shown. **d**, Stacked bar plot of the absolute abundance ($OD_{600}$) of individual biological replicates of the 8-member community over 56 passages (298 generations). **e**, Boxplot showing the distribution of *C. difficile* absolute abundance from the 1$^{st}$ to the 56$^{th}$ passages. Data was based on a long-term passaging experiment shown in **panel d.**

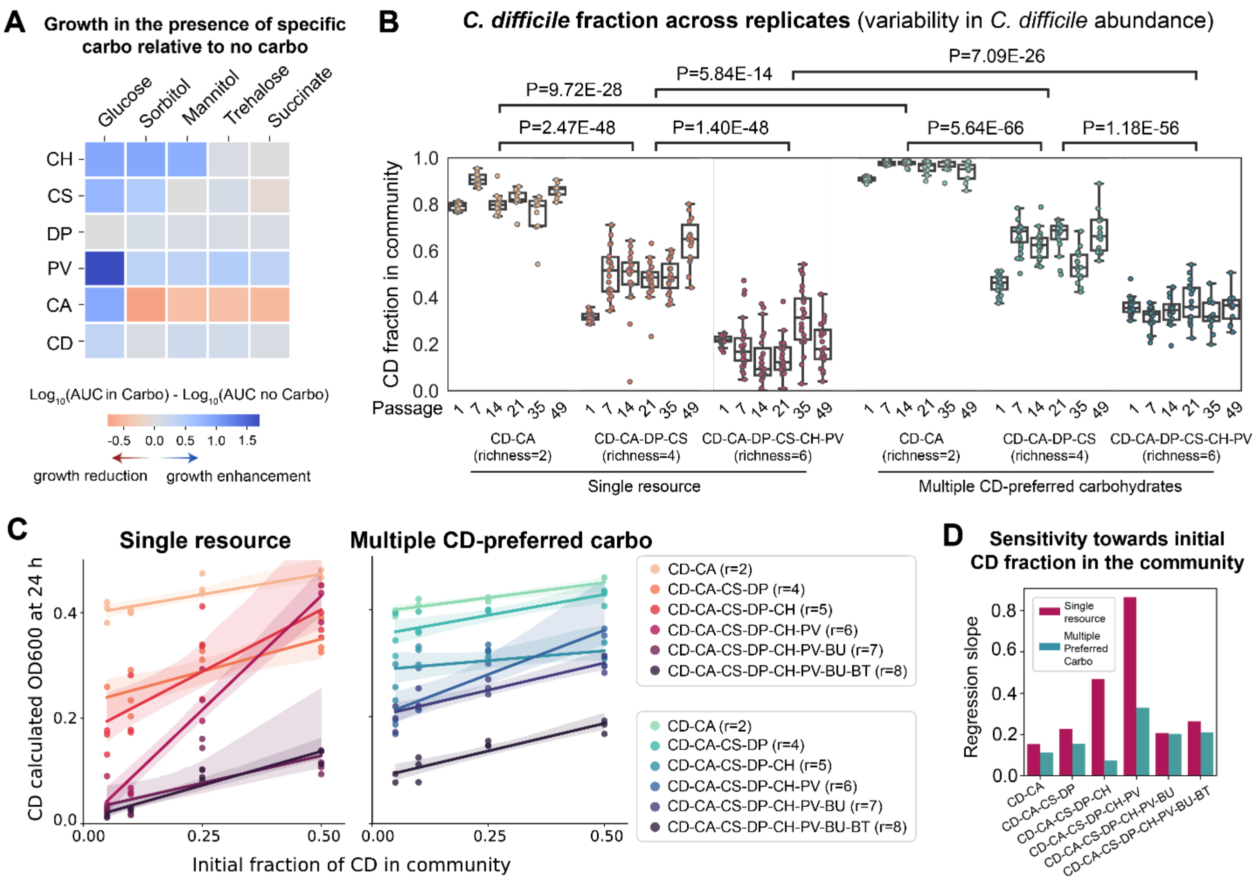


**Supplementary Figure 19. Nutrient environments affect variability in long-term community assembly.** **a**, Heatmap of the Area Under the Curve (AUC) from 24 h monoculture growth data in the presence of specific carbohydrates relative to the absence of carbohydrates. Blue (positive) indicates growth enhancement and red (negative) indicates growth reduction in the presence of specific carbohydrates. **b**, Boxplot of *C. difficile* fraction in the communities across replicates from the 1st to the 49th passages. Plots were based on data from **Fig. 5a**. *p*-values from unpaired *t*-test (two-sided) between different sample groups are shown. **c**, Absolute abundance ($OD_{600}$) of *C. difficile* at 24 h as a function of the initial fraction of *C. difficile* in different synthetic communities with varying species richness. Left panel shows the experiment performed in a media containing a single resource (glucose at 5 g/L), whereas the right panel shows the experiment performed in a media containing multiple *C. difficile*-preferred carbohydrates (glucose, sorbitol, mannitol, trehalose, and succinate at 1g/L each). *C. difficile* was added to communities at 0 h. Datapoints indicate biological replicates (n=3). Lines indicate linear regression with 95% confidence interval (shading). Resident species richness (r) at 0 h is indicated in legend. Calculated $OD_{600}$ is the product of 16S relative abundance and community $OD_{600}$. **d**, Comparison of the regression curve slopes of communities grown in the glucose-only media and the multiple *C. difficile*-preferred carbohydrates media from **panel c**. Larger slope indicates that *C. difficile* final abundance is more affected by variations in the initial abundance values.
